# Genetical and morphological variability of the greater blind mole rat *Spalax microphthalmus* and phylogenetic affinities of large-bodied spalacids (Rodentia, Spalacidae) with a description of *Spalax lyapunovae* sp. nov. from the North Caucasus

**DOI:** 10.1101/2025.10.17.683087

**Authors:** Oleg V. Brandler, Andrey R. Tukhbatullin, Svetlana Y. Kapustina, Fatimat A. Tembotova, Andrey Y. Puzachenko

## Abstract

The greater blind mole rat *S. microphthalmus* is the only species that differs in the number of chromosomes (2n = 60) from the other species (2n = 62) of the genus *Spalax*. It also has the largest range among them. The intraspecific genetic and morphological variability of *S. microphthalmus* has been poorly studied so far. Previously, animals with 2n = 62 were found in the North Caucasus. This was interpreted as a manifestation of intraspecific polymorphism in greater blind mole rats. We investigated the morphometry, as well as the morphological, molecular, and chromosomal variability of *S. microphthalmus* using samples collected from across the entire range of the species. Genetic variability of *S. microphthalmus* was relatively low within the 2n = 60 karyotype range. A comparative, multi-proxy analysis of the species characteristics, as well as phylogenetic analyses within the genus *Spalax* was also performed. Species-level differences were found in all the characteristics studied in the karyotype of the blind mole rat (2n = 62) from the North Caucasus, compared with all the other *Spalax* species, including *S. microphthalmus*. A unique combination of morphological characteristics was described for this blind mole rat, including features that bring it closer to the hypothetical ancestor of all modern species of this genus. A new species is described on this basis: *Spalax lyapunovae* sp. nov. It is necessary to further study the range and the ecology of the new species, which is endemic to the central part of the North Caucasus.

## Introduction

Blind mole rats (subfamily Spalacinae) are highly specialized rodents adapted to an underground lifestyle, inhabiting grassland regions of Eastern Europe and the Eastern Mediterranean (Topachevski 1969). Their exclusively subterranean lifestyle has led to the development in blind mole rats of specific morphological, physiological, and behavioral traits on the one hand, and to lower interspecific morphological variability compared with other rodents, due to the high conservatism of their ecological niche, on the other hand (Nevo 2000).

The systematics of blind mole rats has not been satisfactorily developed to date due to multiple convergent traits limiting the diversity of phenotypes adapted to a burrowing lifestyle (Arslan *et al*. 2016). The division of Spalacinae into two genera, the large-bodied *Spalax* Güldenstaedt, 1770 and the small-bodied *Nannospalax* Palmer, 1903 blind mole rats, is supported by a number of authors (Topachevski 1969; Kryštufek *et al*. 2012; Arslan *et al*. 2016; Németh *et al*. 2024), while some others consider Spalacinae to be monogeneric with a single genus, *Spalax* (Musser & Carleton 2005; Kryštufek & Vohralík 2009). The division of blind mole rats into two genera, initially defined on the basis of morphology, is supported by significant differences in their karyotypic features. All *Spalax* species have similar karyotypes with 2*n* = 62, consisting only of bi-armed chromosomes. The only exception is the greater blind mole rat, *S. microphthalmus* Güldenstaedt, 1770, with 2*n* = 60 (L’apunova *et al*. 1974). In contrast, *Nannospalax* species have extremely variable chromosome sets, represented by 73 chromosomal races with diploid chromosome numbers ranging from 36 to 62, with a diverse combination of acrocentric and bi-armed chromosomes (Arslan *et al*. 2016). Molecular data also support the high differentiation of *Spalax* and *Nannospalax* (Hadid *et al*. 2012; Németh *et al*. 2024).

The greater blind mole rat has the widest geographical range of any *Spalax* species. It covers the area from the Dnieper River in the west to the Volga River and the Ciscaucasia in the east (Topachevski 1969). It is thought to have evolved from the Early-Middle Pleistocene Nogai blind mole rat *S. minor* Topačevski, 1959. Holocene fossils of *S. microphthalmus* have been found in a variety of habitats in localities associated with “krotovina’ loess” (Topachevski 1969). The geographic variability of the greater blind mole rat has not been adequately studied and no subspecies have been identified (Topachevski 1969). Preliminary karyotypic studies have revealed local population variability in the chromosomal rearrangement leading to the formation of a large telocentric chromosome found in the heterozygote with a metacentric chromosome (Puzachenko & Baklushinskaya 1997). A karyotype with 2n = 62 has been described at the southern periphery of the greater blind mole rat distribution in northern Cis-Caucasia (Dzuev & Shogenov 2004; Dzuev *et al*. 2025a). Studies of blind mole rats from this region revealed morphological and physiological features that the authors attributed to adaptation to high altitude habitat conditions (Dzuev & Shogenov 2003, 2004; Dzuev *et al*. 2019; 2025b). The authors were unable to perform a comparative analysis and assess the taxonomic significance of the detected characters due to a lack of data on *S. microphthalmus* from other parts of the range. Dzuev and his colleagues (Dzuev & Shogenov 2003; 2004; Dzuev *et al*. 2019; 2025a) have proposed intraspecific chromosomal polymorphism in the greater blind mole rat. Later, Korobchenko & Zagorodniuk (2009) suggested that the North Caucasian blind mole rats with 2n = 62 belong to giant blind mole rat, *S. giganteus* Nehring, 1898, whose range lies about 100 km to the east, based solely on chromosome numbers. The morphological peculiarities of the North Caucasian blind mole rat were earlier noted by Topachevsky (1969: 216), who suggested that this form may be a distinct subspecies. The taxonomic ambiguity of North Caucasian greater blind mole rats (2n = 62) is one of the unresolved problems of the *Spalax* systematics, which is generally quite stable (Arslan *et al*. 2016; Rusin *et al*. 2024). The intraspecific molecular genetic variability of *S. microphthalmus* has been poorly studied so far. Previous phylogenetic studies of Spalacinae have included a few individuals of *S. microphthalmus*, mainly from the western part of the range or from the southern Don River basin (Hadid *et al*. 2012; Rusin *et al*. 2024; Németh *et al*. 2024).

Clearly, the uncertain species structure and systematic position of the North Caucasian form (2n = 62) remain unresolved issues for the modern *S. microphthalmus*. In order to address these concerns, an examination of greater blind mole rats from all geographical regions of the species’ distribution was conducted to elucidate their intraspecific diversification. A particular emphasis was placed on the characterization of the North Caucasian 62-chromosomal form, with the objective of evaluating its taxonomic status. Karyotype analysis was performed using both standard and differential chromosome banding techniques. Combined phylogenetic analysis of sequences of mitochondrial DNA molecular markers and morphological analysis of craniological characters were performed for *S. microphthalmus* and other species of the genus *Spalax* to reveal intraspecific genetic variability and to determine the position of the North Caucasian form (2n = 62) on the phylogenetic tree.

## Materials and Methods

### Genetic study

#### Karyotyping

Chromosome preparations were obtained from 10 individuals of blind mole rats from 5 localities of the *S. microphthalmus* s. l. range (Table 1 and Fig. 1; Table S1, Supporting information). Of these, 5 individuals (2n = 60) from 4 localities of the *S. microphthalmus* range and 5 individuals (2n = 62, *Spalax* sp.) from the vicinity of Kislovodsk (Stavropol Krai, Russia) in the North Caucasus. The standard technique for preparing chromosome slides from bone marrow was applied (Ford & Hamerton 1956). Routine staining of chromosomes was performed with 2% Giemsa stain. Differential G-banding was made using the method of M. Seabright (1971), and C-banding was carried out according to A.T. Sumner (1972). The stained chromosome slides were observed and photographed using a KEYENCE BZ-9000 microscope with immersion. The karyograms were assembled in Adobe Photoshop v.19.1.2. We followed the chromosome arrangement scheme proposed by L’apunova *et al*. (1974).

**Figure 1.**
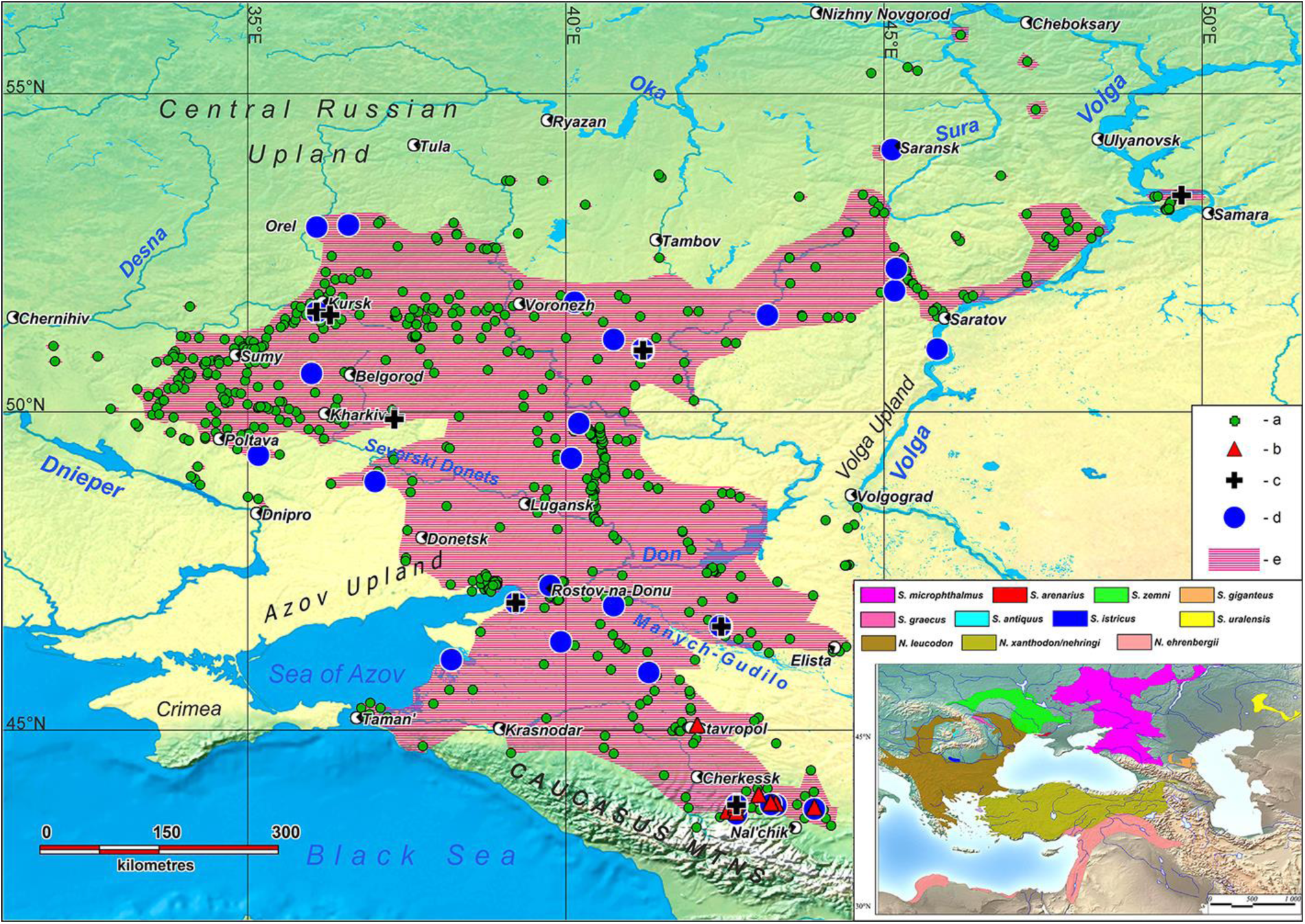
The cadastral map of the localities of *Spalax microphthalmus* s. l. including form with 2*n* = 62 and the sketch map of the species ranges in the subfamily Spalacinae (insert map): *a* – localities of *S. microphthalmus* s. l., *b* – localities of *Spalax* sp. (2n = 62), *c* – localities of the samples of cytogenetic and/or genetic studies, *d* - localities of the samples used in the morphometric study, e – approximated basic range of *S. microphthalmus* s. l.

**Table 1.** The number of blind mole rats of the genus *Spalax* used in the study.

| Sample | Studied by method |  |  |  |  |  |  |
| --- | --- | --- | --- | --- | --- | --- | --- |
|  | Karyology |  |  | Molecular genetic | Morphometric |  |  |
|  | M<sup>1</sup> | F<sup>2</sup> | Total | Total<sup>3</sup> | M<sup>1</sup> | F<sup>2</sup> | Total |
| *Spalax* sp., $2n = 62$ | 4 | 1 | 5 | 6 | 12 | 6 | 18 |
| *S. microphthalmus* | 3 | 2 | 5 | 14(3) | 25 | 18 | 43 |
| *S. zemni* |  |  |  | (5) | 1 | 2 | 3 |
| *S. arenarius* |  |  |  | (3) | 4 | 15 | 18 |
| *S. graecus* |  |  |  | (1) | 16 | 5 | 21 |
| *S. giganteus* |  |  |  | (2) | 17 | 8 | 25 |
| *S. uralensis* |  |  |  |  | 8 | 5 | 13 |
| *S. antiquus* |  |  |  | (3) |  |  |  |
| Total | 7 | 3 | 10 | 34 | 83 | 59 | 142 |

<sup>1</sup>Males; <sup>2</sup>Females; <sup>3</sup>The number of sequences retrieved from GenBank is in brackets.

#### Tissue sampling; DNA extraction, amplification, and sequencing

Tissue samples (kidney, liver, or skin) for the study of the genetic variability of blind mole rats were obtained from the ‘Collection of wildlife tissues for genetic research’ of the Koltzov Institute of Developmental Biology of the Russian Academy of Sciences (CWT IDB), state registration number 3579666. A total of 20 individuals of *S. microphthalmus* s. l. from 11 localities were studied, including six from the North Caucasus (Table 1 and Fig. 1; Table S1, Supporting information). In a previous study, the karyotypes of two individuals from the Kursk region were reported (Puzachenko & Baklushinskaya 1997), and it was established that they had 2*n* = 60 (Table S1, Supporting information). In addition, two specimens of the lesser blind mole rat *Nannospalax leucodon* (Nordmann, 1840) from Odessa Oblast, Ukraine (CWT IDB collection numbers 25248, 25249) were examined for comparative analysis. This study was approved by the Ethics Committee for Animal Research of the Koltzov Institute of Developmental Biology RAS (protocol No 37-25.06.2020).

Genomic DNA was extracted by the standard salt method (Aljanabi & Martinez 1997). A full-length cytochrome *b* (*cytb*) mitochondrial gene was used as a phylogenetic marker. A polymerase chain reaction (PCR) was performed with specific primers SpalaxCBfw2 and SpalaxCBrv2 (Németh *et al*. 2020) in 15 μL volume using 0.3 U HS-Fuzz polymerase and 6 μL 2.5X reaction buffer (Dialat Ltd., Moscow, Russia), 0.3 mM dNTP (Evrogen, Russia), 1pM each primer, 7.85 μL ddH_2_O, and 1 μL DNA 30 ng/μL in a Veriti Thermal Cycler platelet amplifier (Applied Biosystems, Waltham, MA, USA). PCR was performed under the following conditions: initial denaturation for 3 min at 98 °C, followed by 35 cycles of denaturation for 30 sec at 98 °C, 40 sec annealing at 60 °C, 1 min 30 sec extension at 72 °C, and final extension for 7 min at 72 °C. Automated sequencing was performed using a NovaDye Terminator Cycle Sequencing Kit 3.1 (GeneQuest, Moscow, Russia) with the AB 3500 Genetic Analyzer (Applied Biosystems, Waltham, MA, USA), and the Nanofor-05 Genetic Analyzer (Syntol, Moscow, Russia) at the Core Centrum of IDB RAS. Sequence chromatograms were reviewed and manually edited using the SeqMan section of the Lasergene 11 package (DNASTAR, Madison, Wisconsin, USA).

All of the new DNA haplotype sequences were deposited in the NCBI GenBank database under accession numbers PV012717–PV012730.

#### Molecular data analysis

The sequence set obtained was complemented by three *cytb* complete sequences of *S. microphthalmus* that are available on GenBank (NCBI) for the purpose of performing phylogenetic analyses. Sequences of the following species, retrieved from GenBank, were also used in phylogenetic analyses: six other *Spalax* species (Table 1) – *S. antiquus* Méhely, 1909 (3 sequences); *S. arenarius* Reshetnik, 1939 (3); *S. giganteus* (2); *S. graecus* Nehring, 1898 (1); *S. uralensis* Tiflov & Usov 1939 (2); and *S*. *zemni* Erxleben, 1777 (*= S. polonicus* Mehely, 1909) (5), as well as three *Nannospalax* species as an outgroup – *N. ehrenbergi* (3); *N. leucodon* (2); and *N. xanthodon* (3) (Table S2, Supporting information). All the sequences used were aligned using the MUSCLE algorithm (Edgar 2004) and were manually corrected using the MEGA X software package (Kumar *et al*. 2018).

The maximum likelihood (ML) phylogenetic analysis was performed in MEGA X, while Bayesian inference analysis (BI) was performed in MrBayes v3.2.7 (Ronquist *et al*. 2012). Since the *cytb* sequences of *S. uralensis* were 582 bp in length, an ML tree for the short alignment, which included the full set of sequences from all species, was inferred in addition to the full-length (1140 bp) *cytb* alignment that did not include *S. uralensis*.The best nucleotide substitution models were selected through the Bayesian information criterion (BIC) using MEGA X for ML and through the Akaike information criterion (AIC) using jModelTest 2.1.10 (Darriba *et al*. 2012; Guindon & Gascuel 2003) for BI. The TN93+G+I model was used in ML analysis, as well as the GTR+I+G model was used in BI analysis for the full-length alignment. The HKY+G was used for the short alignment in both analyses. BI analysis was based on 2 million generations with each 5000th retained in two MCMC chains with default other settings. The first 25% of the generated trees were discarded as burn-in. Bootstrap analysis with 1000 replicates and posterior probability values were used to estimate branch support for ML and BI trees respectively. Clade support was considered significant when the bootstrap support was 70% (ML) and posterior probability value (BI) was 0.90. The visualization of Bayesian tree was carried out using FigTree v1.4.3 (http://tree.bio.ed.ac.uk). Genetic differences were assessed by pairwise distances (*p*-distance) and Kimura 2-parameter (K2p) in MEGA X. Interspecific genetic distances were calculated for samples larger than 3 individuals.

The evolutionary network of *cytb* haplotypes was constructed using the Median Joining (MJ) method in the HaplowebMaker software (https://eeg-ebe.github.io/HaplowebMaker/ (accessed on 27 January 2025)) (Spӧri & Flot 2020). The genetic variability *cytb* indices including the number of haplotypes (*H*), the haplotype diversity (*Hd*), and the nucleotide diversity (*π*) were estimated for the *S. microphthalmus* (2*n* = 60) sample in DnaSP v.6.12.03 (Rozas *et al*. 2017). The expansion coefficient (*S/k*) was calculated to assess the differences between recent and historical population sizes, as the ratio of the number of variable sites (*S*) to the average number of pairwise nucleotide differences (k) (Peck & Congdon 2004).

### Craniometric study

#### Sample collection and measurements

A total of 142 intact skulls of adult blind mole rats belonging to six species of the genus *Spalax* were analyzed, from 57 localities across the species range. (Table 1 and Fig. 1; Table S1, Supporting information). Among these, the number of skulls belonging to *S. microphthalmus* from 25 localities and *Spalax* sp. (2n = 62) from the 5 localities were 43 and 18, respectively (Fig. 1). A population sample of 122 skulls of adult *S. microphthalmus* (51 females, 60 males) from the “Streletskaya Steppe” (Central Tsernozemny Biosphere Nature Reserve, Kursk district, Kursk oblast, Russia) was used as additional material. Adult specimens were defined by scoring the morphological features of skull structure, such as the development of crests and the chewing surface structure of molars (Topachevski 1969).

Twenty-eight measurements were taken of each skull using digital calipers to the nearest 0.1 mm. The scheme of measurements used is shown in Fig. 2. We used the ratio of a particular measurement to the maximum skull length (MSL) in a number of cases to characterize relative sizes. In addition, the length of incisive foramen and the length and width of upper and lower molars were estimated in 9 specimens of *S. microphthalmus* and 6 specimens of *Spalax* sp. (*2n* = 62). The width and length of the chewing surface of a sample of 23 *Spalax* teeth from the Middle Pleistocene (MIS 9, 11: 337–300, 424–374 ka BP) paleosols of the Otkaznoye locality (44°18’56.97’’N, 43°50’44.29’’E, Stavropol Krai, Russia (Markova 2006; Bolikhovskaya *et al*. 2016)) were also measured: 3 M1, 4 M2, 4 M3, 6 m1, 4 m2, and 2 m3. We used the dental terminology of upper and lower molars according to (Topaczewski, 1969; Sarika and Sen, 2003; Loṕez-Antonyanzas, 2012; Erten, 2018), which are shown on molars of *S. microphthalmus* juveniles in Fig. 3.

**Figure 2.**
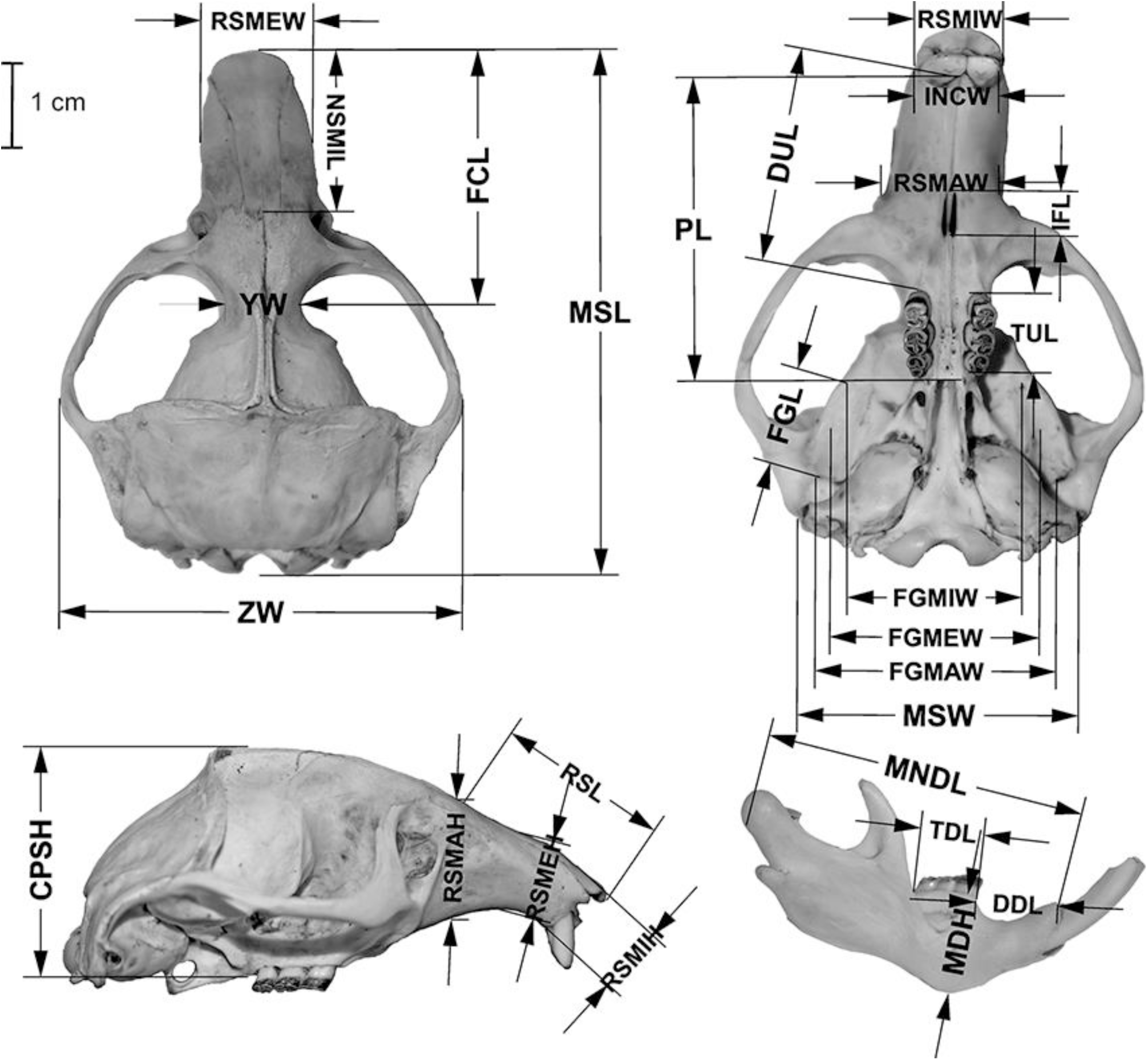
The scheme of blind mole rat skull measurements used in the study (skull of *Spalax microphthalmus*): MSL – maximum skull length, NSMIL – minimal length of nasal bone, FCL – facial length, PL – palatal length, RSL – rostral length, DUL – length of upper diastema, TUL – alveolar length of upper tooth-row, IFL – length of *foramen incisivum*, FGL – length of *fossa glenoidea* (glenoid cavity), ZW – zygomatic width, YW – interorbital width, MSW – mastoid width, RSMIW, RSMEW, RSMAW – minimal, intermediate and maximal width of rostrum, INCW – double upper incisive width, FGMIW, FGMEW, FGMAW – minimal, intermediate and maximal width of skull base between the left and right *fossa glenoidea*, RSMIH, RSMEH, RSMAH – minimal, intermediate and maximal height of rostrum, CPSH – skull height, MNDL – condylar length of mandible, DDL – length of lower diastema, TDL – alveolar length of lower tooth-row, MNDH – height of horizontal branch of mandible, and, in addition, CNDL, CNDW – the length and width of the condyle (not shown).

**Figure 3.**
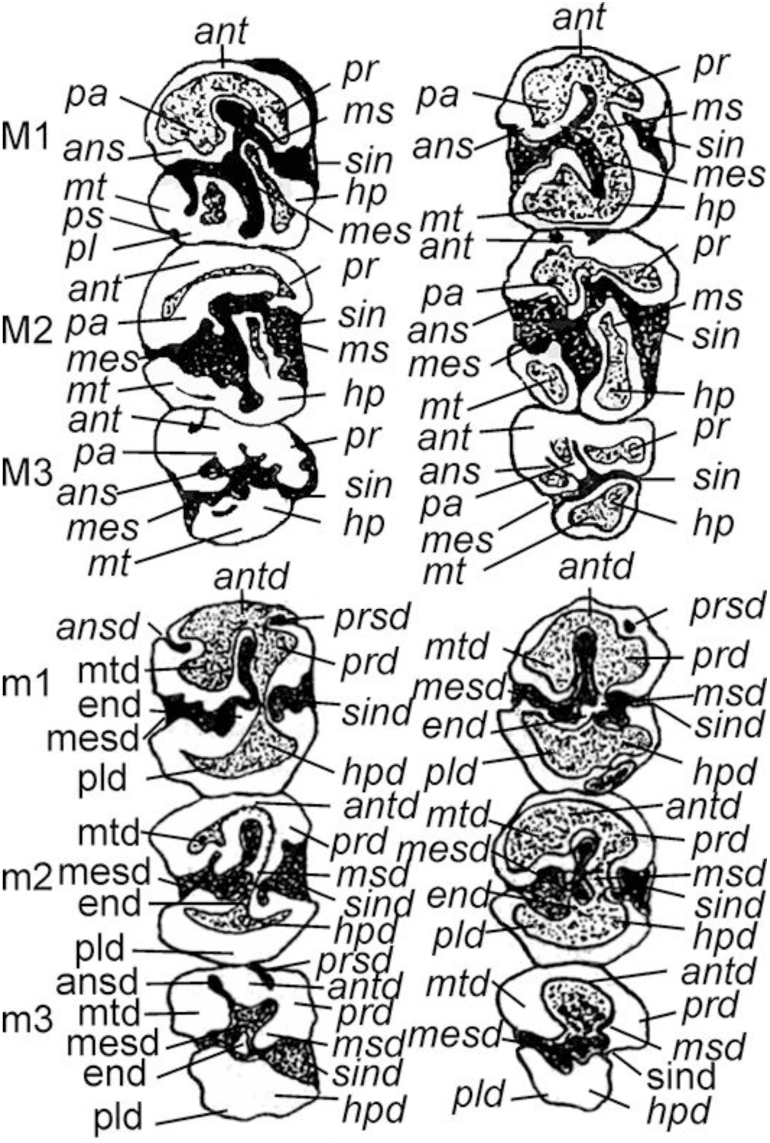
Occlusal surfaces of upper (M1–M3) and lower (m1–m3) molars of *S. microphthalmus* juveniles (modified from (Puzachenko 1991)) and dental terminology used in the study (compiled by (Topachevski 1969; Sarica & Sen 2003; Loṕez-Antoñanzas 2012; Erten 2018)): 1) upper molars: *ant* – anterocone (anteroloph), *pa* – paracon (protoloph), *pr* - protocon, *ms* – mesocon (mesoloph), *mt* – metacone (mtaloph-posterioloph), *hp* – hypocon, *pl* - posteroloph, *ans* - anterosinus, *mes* - mesosinus, *ps* – posterosinus; 2) lower molars: *antd* – anteroconid (anterolophid), *mtd* – metaconid (metalophid), *msd* – mesoconid (mesolophid), *end* – entoconid (hypolophid), *prd* - protoconid, *hpd* - hypoconid, *pld* – posterolophid, *ansd* – anterosinisid, *prsd* – protosinusid, *sind* - sinusid, *mesd* – mesosinusid.

Common methods of univariate statistical analysis used include descriptive statistics (median, mean, standard error of the mean, and 95% confidence limits for the mean and percentiles), Pearson’s correlation, ANOVA, and the nonparametric Mann-Whitney U test. Two *morphospaces* (Puzachenko 2023) were developed using the nonmetric multidimensional scaling (NMDS (Kruskal 1964, Davison 1983)) based on the matrices of Euclidean distances between all pairs of skulls (general skull size model, SZM) and Kendall’s tau-b (corrected for ties) associations (Kendall 1938) (general skull shape model, SHM). The structure of the SZM and SHM models was studied using correlation analysis and analysis of variance (ANOVA). For the species samples, the medians of the coordinates of the SZM and SHM models were calculated. In the text, the NMDS axes were denoted as E1, E2, etc. for SZM models, and as K1, K2, etc. for SHM models. We used components of variance analysis (Solomon 2005) to estimate the *a priori* taxonomy effect on the model axis variations. The species sample centroid coordinates were used as variables in a cluster analysis.

Statistical analysis was carried out using STATISTICA v. 8.0 (StatSoft, Tulsa, Oklahoma), PAST v. 3.12 (Hammer & Harper 2007), NCSS v. 12 Statistical Software (ncss.com/software/ncss), and Dendroscope ver. 3.8.3 (Huson & Scornavacca 2012).

## Results

### Karyotype variability in S. microphthalmus s. l

Of the ten *S. microphthalmus* s. l. individuals examined, five had chromosome sets with 2n = 60, NFa = 116, and NF = 120 (Fig. 4). The provenance of the specimens under consideration were the *terra typica* of the species in the Novokhopersk steppes (Topachevski 1969), as well as the Kursk region, the southern bank of the Don River delta, and the northern bank of the Manych-Gudilo channel (Table S1, Supporting information). This karyotype was found to be similar to that which had previously been described for *S. microphthalmus* (L’apunova *et al*. 1974). The other five blind mole rats were from the North Caucasus and had 2n = 62, NFa = 120, and NF = 124 (Fig. 5A). This chromosome set consisted of five pairs of medium and small metacentrics, 12 pairs of submetacentrics decreasing in size, and 13 pairs of subtelocentrics, including two pairs of the largest elements of the set. The X chromosome was a large metacentric or submetacentric, and the Y chromosome was a small subtelocentric. This karyotype corresponds to the one previously described in the Northern Caucasus (Dzuev & Shogenov 2004; Dzuev *et al*. 2025a), with the exception of the ratio of metacentrics and submetacentrics. This discrepancy could be explained by the ambiguity in the morphology of small bi-armed chromosomes with different levels of spiralization.

**Figure 4.**
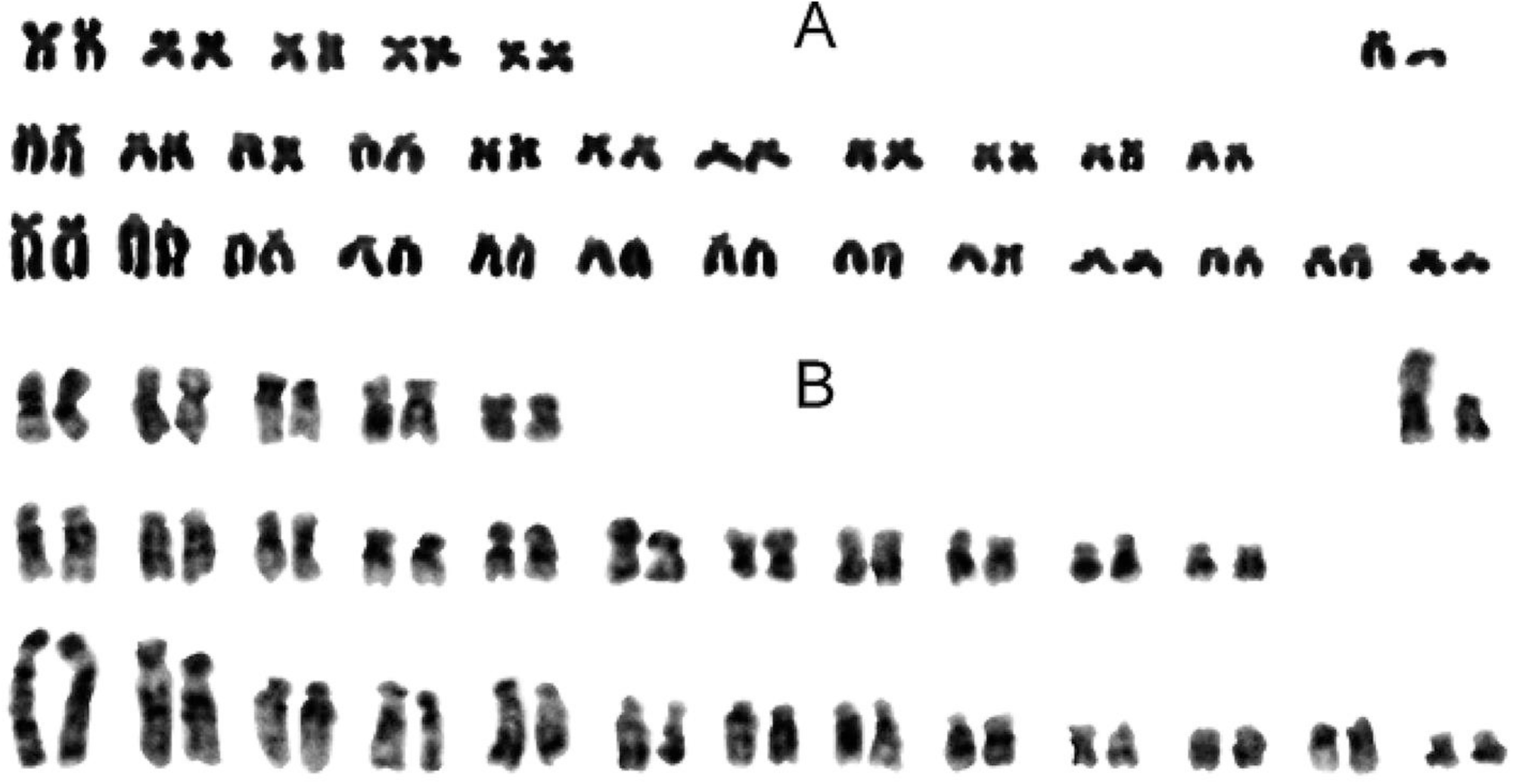
Karyotype of the *S. microphthalmus* (2n = 60) male with routine painting (A) and C-banding (B) of chromosomes.

**Figure 5.**
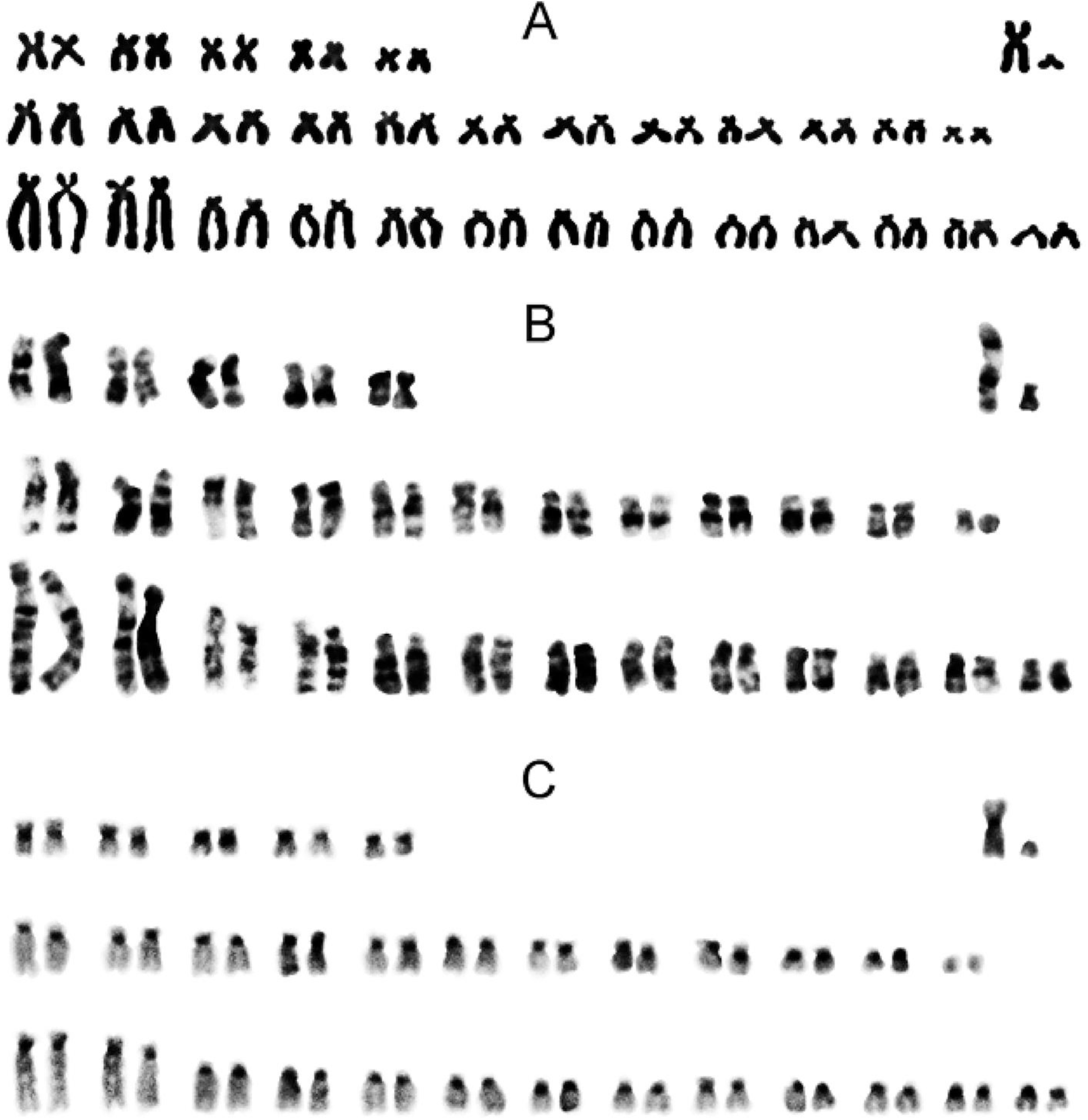
Karyotype of the *Spalax* sp. (2n = 62) male with routine painting (A), G-banding and C-banding (C) of chromosomes.

The presence of heterochromatin blocks was detected on all chromosomes of the 62-chromosome karyotype in the centromeric regions, with the exception of the smallest pair of submetacentrics (Fig. 5C). The X chromosome had a large subcentromeric block, and the Y chromosome was almost entirely heterochromatized. This result was consistent with the textual description of C-painting provided in the publication of Dzuev and Shogenov (2004): “…pericentromeric heterochromatin blocks have been detected in nearly all chromosomes…”. The near-centromeric heterochromatin blocks were detected on all chromosomes in the normal karyotype of *S. microphthalmus* (2n=60) and were consistent with the previously published C-banded karyotype (Puzachenko & Baklushinskaya 1997).

### Cytb variability in the genus Spalax

In a total sample of 23 complete *cytb* sequences (1140 bp) from *S. microphthalmus* s. l., thirteen distinct haplotypes were identified in *S. microphthalmus* (2n = 60) and two haplotypes in *Spalax* sp. (2n = 62). The 60 and 62 chromosome forms did not share any common haplotypes. The mean nucleotide composition was as follows: A=31%, T=32%, G=13%, and C=24%. The dataset contained 106 variable positions, 96 of which were parsimony informative. The total number of mutations identified was 107, of which 18 were non-synonymous substitutions. Within the cytochrome *b* protein sequence, which contains 379 amino acids, 11 were found to be variable in the total sample of *S. microphthalmus* s. l. Six of these variable amino acids distinguished the 60- and 62-chromosome forms.

The topologies of the phylogenetic trees inferred from maximum likelihood (ML) and Bayesian analysis (BI) were found to be similar (Fig. 6; Fig. S1, Supporting information). All the studied *Spalax* species formed separate branches, a finding that received a high level of statistical support. However, the position of the *S. giganteus* cluster, as well as the combined *S. giganteus*/*S. uralensis* cluster (Fig. 6A), lacked statistical support across all trees. The well-differentiated branches of *Spalax* sp. (2n = 62) and *S. microphthalmus* (2n = 60) grouped together into a common clade in all analyses, albeit with low bootstrap values.

**Figure 6.**
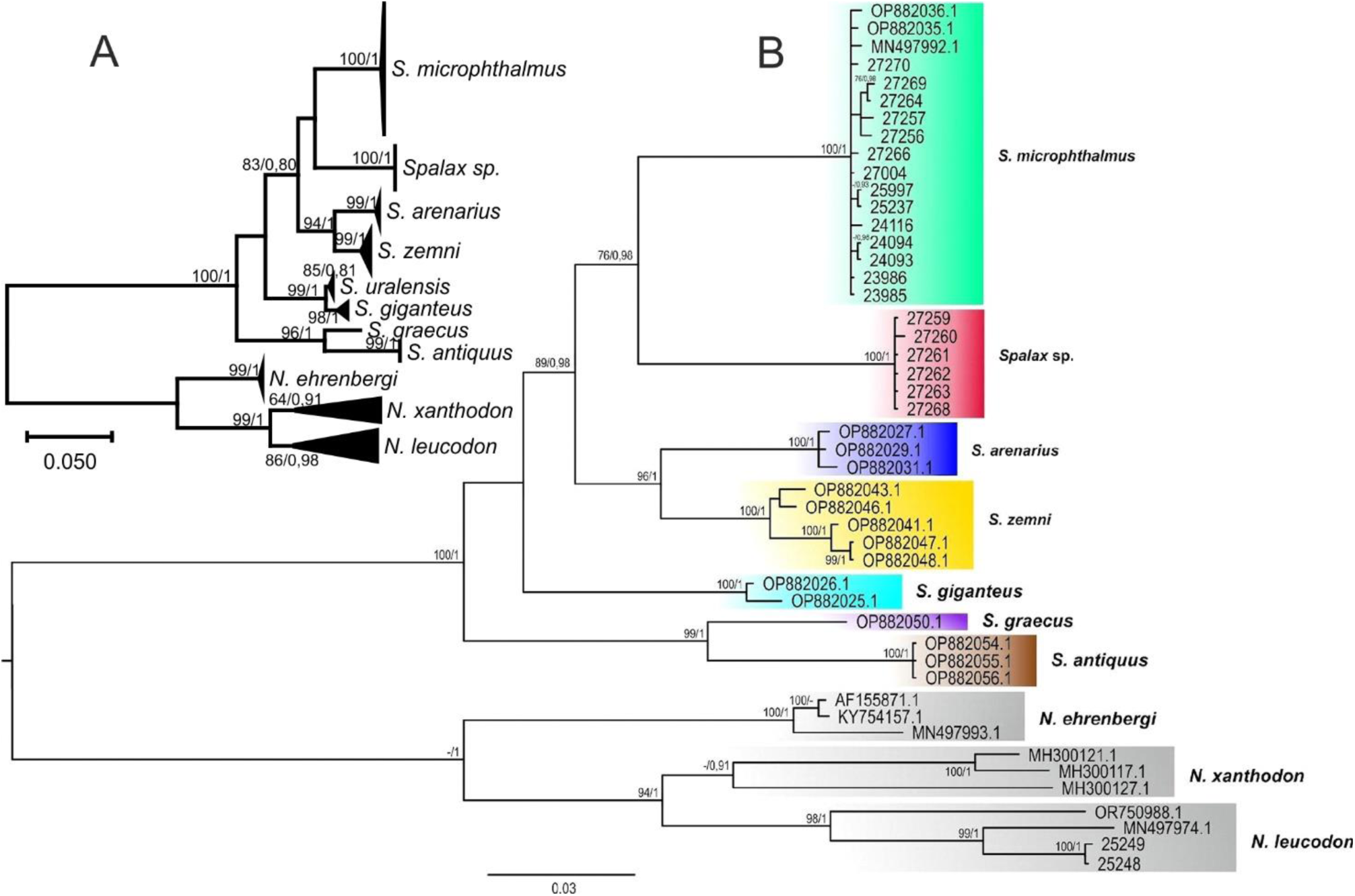
Phylogenetic trees of blind mole rats based on the mitochondrial *cytb* gene: (A) ML tree for the short alignment (582 bp); (B) BI tree for the full-length alignment (1140 bp) not including *S. uralensis*. The values at the nodes represent bootstrap support for the corresponding nodes in the ML tree and posterior probability after slash. Values below 70% (ML) and 0.90 (BI), as well as for minor nodes, are not shown.

In constructing the median network of haplotypes, a direct link was identified between the evolutionary lineage of the *Spalax* sp. haplotypes and one of the haplotypes (c13 in the specimen 27257) found at the southern frontier of the *S. microphthalmus* (2n = 60) range (Table S1 and Fig. S2, Supporting information). The analysis of the sequence of the c13 haplotype revealed that its position on the network was determined by two substitutions. It was found that the c13 haplotype possessed nucleotide A at the locus 495, which is species-specific for *S. microphthalmus* (2n = 60), as well as G at the locus 813, which is specific to *Spalax* sp. (2n = 62). This was in contrast to the other haplotypes of this lineage (c10-c12), which were also distributed in southern populations. The exclusion of this haplotype from the analysis resulted in the construction of a network (Fig. 7) on which the *Spalax* sp. lineage evolved from the central group of hypothetical haplotypes in the same way as most of the other *Spalax* species. The characteristics of the key substitutions, in conjunction with the position of the sample 27257 haplotype on the phylogram (Fig. 6B), indicate a more probable convergent origin of these substitutions in haplotype c13 than homologies with the *Spalax* sp. haplotypes. In this case, the network structure suggested the origin of most *Spalax* species, including *S. microphthalmus* (2n = 60), *Spalax* sp. (2n = 62), and *S. giganteus*, as the result of a radial divergence from a common ancestor followed by independent evolution.

**Figure 7.**
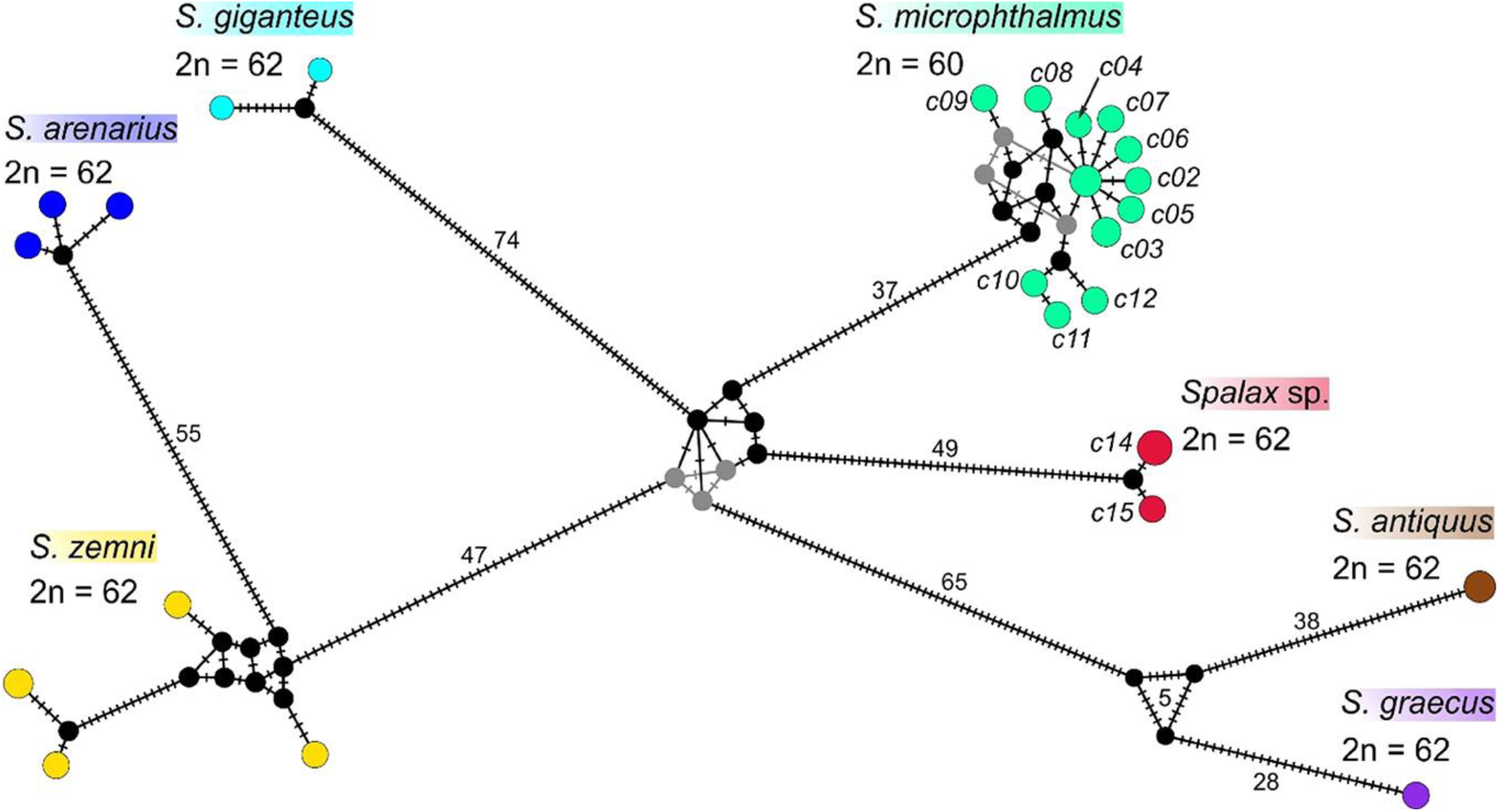
Evolutionary *cytb* haplotype networks of *Spalax*. Haplotype labels of *S. microphthalmus* and *Spalax* sp. correspond to Table S1, Supporting information. Haplotype *c13* was excluded. The colors correspond to Fig. 6B.

The average genetic distances between the blind mole rat species studied are presented in Table 2. All the *Spalax* species exhibited significant genetic differentiation. The genetic distances between *Spalax* sp. (2n = 62) and *S. microphthalmus* (2n = 60) reached notional interspecific values (>5%) for mammals (Baker & Bradley 2006), which were commensurate with the differences observed between other *Spalax* species, including both of these species and *S. giganteus*. The wide-ranging species *S. microphthalmus* (2n = 60) showed remarkably low intraspecific genetic variability, with intraspecific genetic distances based on the *cytb* gene being comparable with those observed in closely related species with significantly smaller habitat ranges, such as *S. arenarius*, *S. graecus*, and *S. antiquus* (Rusin *et al*. 2024). This result was consistent with the low level of *S. microphthalmus* genetic variability of the control region of mtDNA, which was found earlier (Matveeva *et al*. 2019). The genetic variability indices of the *S. microphthalmus* (2n = 60) *cytb* sample (N = 17) were: *H* – 13; *H_d_* – 0.963; *π* – 0.0028; *S* – 20; *k* – 3.191. The expansion coefficient (*S/k*) was 6.268.

**Table 2.** The mean genetic distances between the examined blind mole rat species. The *p*-distance values are below the diagonal, the *K2p* values are above the diagonal, and the intraspecific *p*-distances for samples larger than 3 individuals are on the diagonal

| Species | *S. sp.* (2n = 62) | *S. microphthalmus* | *S. giganteus* | *S. arenarius* | *S. zemni* | *S. graecus* | *S. antiquus* | *N. ehrenbergi* | *N. leucodon* | *N. xanthodon* |
| --- | --- | --- | --- | --- | --- | --- | --- | --- | --- | --- |
| *S. sp.* (2n = 62) | **0.0006** | 0.0876 | 0.1021 | 0.0905 | 0.1014 | 0.1206 | 0.1314 | 0.2002 | 0.2118 | 0.2028 |
| *S. microphthalmus* | 0.0812 | **0.0028** | 0.0914 | 0.0905 | 0.0945 | 0.1220 | 0.1212 | 0.2034 | 0.2150 | 0.2027 |
| *S. giganteus* | 0.0932 | 0.0844 | - | 0.1021 | 0.0951 | 0.1120 | 0.1179 | 0.1972 | 0.2004 | 0.1910 |
| *S. arenarius* | 0.0833 | 0.0837 | 0.0930 | - | 0.0617 | 0.1023 | 0.1156 | 0.1937 | 0.2073 | 0.2045 |
| *S. zemni* | 0.0925 | 0.0869 | 0.0872 | 0.0583 | **0.0132** | 0.1025 | 0.1147 | 0.1975 | 0.1986 | 0.1989 |
| *S. graecus* | 0.1088 | 0.1101 | 0.1018 | 0.0939 | 0.0942 | - | 0.0660 | 0.1963 | 0.2160 | 0.1988 |
| *S. antiquus* | 0.1175 | 0.1095 | 0.1066 | 0.1050 | 0.1042 | 0.0623 | - | 0.2084 | 0.2133 | 0.2010 |
| *N. ehrenbergi* | 0.1729 | 0.1757 | 0.1708 | 0.1681 | 0.1712 | 0.1705 | 0.1792 | - | 0.1441 | 0.1347 |
| *N. leucodon* | 0.1820 | 0.1846 | 0.1738 | 0.1786 | 0.1725 | 0.1851 | 0.1831 | 0.1286 | **0.0181** | 0.1241 |
| *N. xanthodon* | 0.1754 | 0.1755 | 0.1664 | 0.1764 | 0.1725 | 0.1728 | 0.1743 | 0.1212 | 0.1124 | - |

### Craniometrical variability in the genus Spalax

The SZM and SHM models had four and three coordinates respectively (Table S3, Supporting information). The relative components of variance of coordinates E1 and K1 due to the conventional taxonomy in the genus *Spalax* were 83.7% and 80.4% respectively. The next most important coordinates that show a “taxonomic signal” were K2, E4, and K3. The proportion of explained variance attributed to the E1–E4 coordinates varied from 78% (DDL) to 97% (MSL); on average, it was 90%. Thus, the model satisfactorily describes variability in skull size. The high mean proportion of variance (0.81) of the skull measurements attributed to coordinate E1 indicates the dominance of skull variability associated with the overall skull size.

The indicator measurements of size variation were MSL, RSMIH, RSMEW, PL, MSW, FGMAW, INCW, MNDL, and others (Fig. 8A; Table S3 and Figs. S3A and S4A, Supporting information). Interorbital width (YW) correlated with both the E2 (*r* = 0.67) and E3 (*r* = 0.65) coordinates. The projections of the species sample centroids onto coordinates E1 and E2, arranged along E1 according to the disparities in species-specific overall skull size. In this sequence, the skull of *Spalax* sp. (2n = 62) occupies the outermost position as the smallest on average. At the opposite end of the row were the largest members of the genus *Spalax*: *S. uralensis* and *S. giganteus*. Due to the fact that the sample of *S. arenarius* consisted mainly of female skulls, in Fig. 8A, the centroid of this species is located in close proximity to the centroid of the small *Spalax* sp. (2n =62).

**Figure 8.**
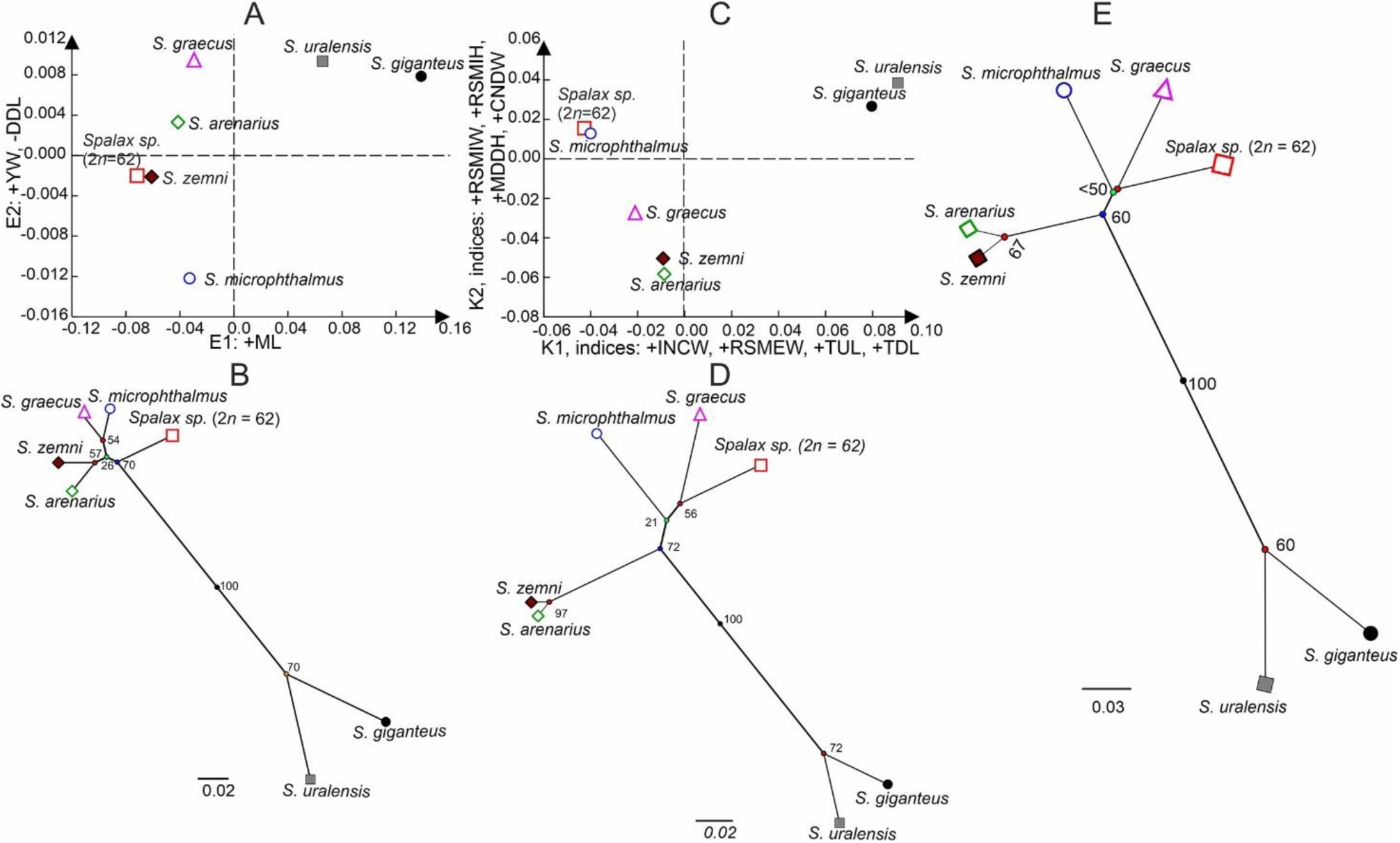
The projections of the species sample centroids onto the E1, E2, (A) and K1xK2 coordinates of the SZM and SHM models, respectively, and the radial classification trees (Euclid distances, UPGMA method) of the species centroids based on the E1 – E4 coordinates of the SZM model (B) and K1 – K3 coordinates of the SHM model (D). E – Radial classification trees (Euclid distance, UPGMA method) of the species centroids based on PC1 – PC4, which accounted for 99.8 % of the E1 – E4 and K1 – K3 variance. The + or - sign in front of the measurement abbreviations corresponds to the sign of the correlation coefficient between the variable and the morphospace coordinate. The numbers near the tree nodes estimate the bootstrap support as a percentage of 1000 iterations.

Coordinate K1 of the SHM showed a weak but significant correlation with the MSL (*r* = 0.56). This indicates the presence of an allometric pattern, i.e. the shape of a skull depends partly on its general size. K1 most strongly correlates with indices TUL/MSL (*r* = 0.80), TDL/MSL (*r* = 0.73), INCW/MSL (*r* = 0.79), and RSMEW/MSL (*r* = 0.77), i.e., with the relative sizes of tooth rows, the width of the upper incisors, and rostrum width (Fig. 8C; Table S3 and Figs. S3B and S4C, Supporting information). The K2 coordinate demonstrates the strongest correlation with indices RSMIW/MSL (r = 0.61), RSMIH/MSL (r = 0.54), MNDH/MSL (r = 0.60), CNDW/MSL (*r* = 0.58), and YW/MSL (*r* = -0.57).

Cluster analysis of sample centroids over all coordinates of the SZM or SHM showed a strong separation of *S. uralensis* and *S. giganteus* (Puzachenko 1993) from all the other species of the genus by size and shape of the skull (Fig. 8B and 8D, respectively). This finding is consistent with the placement of these species on a distinct branch of the phylogenetic tree (Fig. 6A). *Spalax* sp. (2n = 62), as the smallest representative of the genus, occupied a near basal position in the classification tree for all the other species (Fig. 8B). It was closest to *S. graecus*, then to *S. microphthalmus*, and was well differentiated from both *S. zemni* and *S. arenarius* on the tree reproducing similarity in overall skull shape between the taxa (Fig. 8D). At last, we applied PCA of the coordinates of the SZM and SHM together to combine information from both models (Fig. 8E; Fig. S5 and Table S4, Supporting information). In accordance with this synthesis, *Spalax* sp. (2n = 62) shares a “subclade” together with *S. graecus* and *S. microphthalmus*.

In general, the results of the craniometric variability study agree with the results of genetic studies and do not contradict the hypothesis of radial speciation of the modern *Spalax*. Also, a certain convergence of *Spalax* sp. (2n = 62) with *S. microphthalmus* and *S. graecus* does not contradict the evolutionary *cytb* haplotype networks of *Spalax* (Fig. 7).

### A morphological comparison of the North Caucasian Spalax sp. (2n = 62) with members of the genus Spalax

The North Caucasian blind mole rat is a typical member of the genus *Spalax*, based on a combination of the following craniometric characteristics (Topachevski 1969): the supracondylar foramen is always absent; pharyngeal tubercles (*tuberculum pharingeum laterale*) are narrow, elongated as narrow ridges on the edges of the basisphenoid bone (*corpus ossis sphenoidalis*) and the basilar part of the occipital bone; the basisphenoid bone and the basilar part of the occipital bone are approximately level with each other and the skull base is not “fractured”; the corono-alveolar notch (incisura corono-alveolaris) is strongly developed and sharp; the fossa between the corono-alveolar and corono-condilar notches is deep; and the angular process of the mandible is attached to the alveolar process (see SM 2 for more detail).

*Spalax* sp. (2n = 62) differs from all the other recent species of *Spalax* primarily by its smaller skull size. Nevertheless, there is no morphometric hiatus between most skull measurements of this form and those of species other than *S. uralensis* and *S. giganteus* (Table S5, Supporting information). The clearest differences (at *p*<0.0001) between *Spalax* sp. (2n = 62) and all the other species were found for MSL, ZW, CPSH, INCW, TUL, TDL, and FGL. At the same time, *Spalax* sp. does not differ from *S. graecus* and *S. microphthalmus* on the RSMIW and RSMIH. There were no significant differences in the RSMAH between *Spalax* sp. (2n = 62) and *S. graecus*, *S. microphthalmus*, and *S. arenarius*. The FGMIW in *Spalax* sp. (2n = 62) is approximately equal to that of *S. microphthalmus* and *S. arenarius*. The interorbital width of the skull is characterized by low variability within the genus *Spalax*. The mean value of YW/MSL varies between 11.9% (*S. giganteus*) and 14.9% (*S. graecus*) in adults. Regarding the relative size of the YW, *Spalax* sp. (2n = 62) is indistinguishable from all the other species with the widest (15.5%) interorbital part of the skull. The variation of some other indices demonstrated the unique characteristics of *Spalax* sp. (2n = 62) (Fig. S6, Supporting information). The latter has, on average, the highest rostrum, the longest nasal bones (length along the suture between the left and right bones), the widest skull base (MSW), the longest mandible (MNDL) and the shortest *fossa glenoidea* (FGL). According to our data, *Spalax* sp. (*2*n = 62) has a considerably shorter *foramen incisivum* (3.47), at least in comparison with *S. microphthalmus* (5.14) (Table 3), or perhaps with the other *Spalax* species. The average length of the IFL is 18.7% of the length of the upper diastema in *Spalax* sp. (2n = 62) and 24.2% in *S. microphthalmus*. Therefore, we consider the length of the *foramen incisivum* to be a good characteristic to distinguish the North Caucasian form from *S. microphthalmus*. It is surprising that the relative length of IFL in *Spalax* sp. (2n = 62) corresponds to this parameter, which is characteristic of species of the genus *Nannospalax* Palmer, 1903 (*N. ehrenbergi* (Nehring, 1898) (Hamad *et al*. 2016)).

**Table 3.** A preliminary comparison of *Spalax* sp. (2n = 62) and *S. microphthalmus* in length of *foramen incisivum* (IFL, *N* = 18, 9), length of longitudinal section of upper incisor at the level of upper incisor alveoli (INCUL, *N* = 6, 9), and lengths of transverse and longitudinal sections at the level of lower incisor alveoli (INCDW, INCDL, *N* = 6, 9), length (LM1–LM3, Lm1–Lm3, *N* = 6, 9) and width of molars (WM1–WM3, Wm1–Wm3, *N* = 6, 9) and their indexes (%). M-U, Z – result of the Mann-Whitney U Test of a hypothesis of equality of means (*p*)

| VAR | *Spalax* sp. (2n = 62) |  |  | *S. microphthalmus* |  |  | M-U, Z | *p* |
| --- | --- | --- | --- | --- | --- | --- | --- | --- |
|  | M±m | Med | Min-Max | M±m | Med | Min-Max |  |  |
| IFL | 3.47±0.12 | 3.43 | 2.20–4.54 | 5.35±0.15 | 5.3 | 4.73–5.95 | -4.14 | <0.0001 |
| IFL/DUL | 18.7±0.75 | 18.3 | 12.8–24.4 | 24.2±0.78 | 24.3 | 20.1–27.6 | -3.45 | 0.0002 |
| INCUL | 2.84±0.073 | 2.87 | 2.63–3.07 | 3.09±0.134 | 3.03 | 2.69–3.95 | -1.30 | n.s. |
| INCW/INCUL | 134.3±3.5 | 137.3 | 122.8–142.7 | 138.9±3.7 | 139.4 | 115.9–153.9 | -0.7 | n.s. |
| INCDW | 3.41±0.091 | 3.35 | 3.18–3.77 | 3.78±0.106 | 3.7 | 3.41–4.38 | -2.18 | 0.026 |
| INCDL | 3.43±0.09 | 3.40 | 3.11–3.71 | 3.60±0.095 | 3.68 | 3.13–3.94 | -1.24 | n.s. |
| INCDW/INCDL | 99.5±1.9 | 99.8 | 92.9–104.4 | 105.1±2.1 | 105.1 | 92.7–144.3 | -1.65 | n.s. |
| LM1 | 2.68±0.07 | 2.72 | 2.38–2.88 | 2.9±0.08 | 3.02 | 2.46–3.24 | -1.06 | n.s. |
| WM1 | 2.65±0.02 | 2.67 | 2.59–2.69 | 2.67±0.03 | 2.69 | 2.52–2.77 | -0.65 | n.s. |
| WM1/ LM1 | 99.2±3.01 | 97.5 | 92.3–113.3 | 92.7±3.25 | 91.9 | 82.21–106.71 | 0.38 | n.s. |
| LM2 | 2.43±0.09 | 2.44 | 2.11–2.67 | 2.47±0.07 | 2.42 | 2.66–2.85 | -0.29 | n.s. |
| WM2 | 2.61±0.04 | 2.63 | 2.47–2.72 | 2.66±0.05 | 2.72 | 2.42–2.89 | 0.38 | n.s. |
| WM2/ LM2 | 108.8±5.61 | 108.1 | 93.8–128.9 | 107.9±2.13 | 110.8 | 97.9–115.1 | -0.05 | n.s. |
| LM3 | 2.19±0.09 | 2.21 | 1.78–2.41 | 2.10±0.06 | 2.14 | 1.65–2.25 | 1.17 | n.s. |
| WM3 | 2.23±0.03 | 2.22 | 2.09–2.34 | 2.23±0.04 | 2.19 | 2.06–2.41 | 0.23 | n.s. |
| WM3/ LM3 | 102.9±5.86 | 97.2 | 93.1–131.2 | 106.9±3.8 | 104.4 | 97.5–134.0 | -1.47 | n.s. |
| Lm1 | 2.45±0.06 | 2.46 | 2.22–2.59 | 2.64±0.05 | 2.71 | 2.28–2.74 | -2.30 | 0.017 |
| Wm1 | 2.46±0.04 | 2.45 | 2.31–2.58 | 2.51±0.03 | 2.54 | 2.39–2.64 | -0.65 | n.s. |
| Wm1/ Lm1 | 100.5±1.47 | 100.9 | 94.6–103.9 | 95.3±1.45 | 95.1 | 88.8–105.1 | 1.94 | 0.05 |
| Lm2 | 2.38±0.05 | 2.41 | 2.21–2.54 | 2.48±0.05 | 2.50 | 2.22–2.78 | -1.24 | n.s. |
| Wm2 | 2.40±0.04 | 2.39 | 2.30–2.55 | 2.59±0.04 | 2.55 | 2.46–2.77 | -2.53 | 0.01 |
| Wm2/ Lm2 | 100.7±1.77 | 101.9 | 94.5–105.6 | 105.2±1.66 | 102.4 | 99.8–113.7 | -1.0 | n.s. |
| Lm3 | 2.07±0.08 | 2.10 | 1.72–2.29 | 2.40±0.09 | 2.39 | 1.94–2.71 | -2.06 | 0.04 |
| Wm3 | 2.18±0.05 | 2.17 | 1.84–2.53 | 2.28±0.07 | 2.29 | 1.84–2.53 | -1.35 | n.s. |
| Wm3/ Lm3 | 105.6±3.19 | 101.1 | 96.3–118.0 | 95.2±2.02 | 94.6 | 85.4–107.5 | 2.42 | 0.01 |

The relatively long mandible (the measurement includes the length of the condylar process, indirectly) probably compensates the short length of the glenoid cavity in *Spalax* sp. (2n = 62). On average, a diastema in this form is about 25.2% (21.9-30.5%) of the mandible length and 15.6% (13.6-17.4%) of the skull length. This value is less than that of the *S. microphthalmus* (27.0%, 22.8-30%; and 16.4%, 13.3-18.8%, respectively) or the *S. arenarius* (27.4%, 25.5-30.1%; and 16.3%, 14.6-18.1%, respectively) and corresponds to the relative diastema length in the *S. graecus* (25.3%, 22.6-28.3%; and 14.98%, 13.4-17.4%, respectively). It should be noted that the smallest relative length of the lower diastema among *Spalax* was found in *S. giganteus* and *S. uralensis* (14.6% and 14.76%, respectively). The relative length of the TDL is about 22.6% (19.6-27.8%) of the mandible length that is approximately the same as in the *S. microphthalmus* (22.5%, 18.8-26.0%) and less than that of the *S. graecus* (24.5%, 22.3-27.2%). Therefore, it is more likely that the relative elongation of the mandible in *Spalax* sp. (2n = 62) is due to the longer condylar process itself rather than the diastema or horizontal branch of the mandible.

In *Spalax* sp. (2n = 62), short glenoid cavities, combined with the relatively small difference in distance between their anterior and posterior edges (FGMIW and FGMAW, respectively), result in a rather sharp angle between the imaginary lines drawn along the surfaces of the glenoid cavities (Fig. S7A, Supporting information). A rough estimate of the angle between the glenoid cavities using only the FGMIW, FGMAW, and FGL measurements in *Spalax* sp. (2n = 62) was approximately 18.8°, while in other species, this value exceeded 19° (19.7° to 23.8°). In addition, this form had a relatively narrow articular head (length/width ratio 28.5% (24.4-33.8%), which should slightly limit the possibility of lateral movements of the mandible. In comparison, the index was 38% (28.1-47.3%) in *S. giganteus*, 30.1% (24.5-41.4%) in *S. graecus*, and 31.1% (25.1-36.9%) in *S. microphthalmus*.

In *Spalax* sp. (2n =62), the position of the upper alveolar ridge, which is not strongly developed, relative to the first upper molar differs markedly from that of *S. microphthalmus* (the alveolar ridge is located close to M1), but it is similar to that of the other members of the genus (Fig. 9). *Spalax* sp. (2n = 62) probably differs from the other species by the shape of the posterior palatine notch. A characteristic feature of *Spalax* sp. (2n =62) is the convex shape of the posterior margin of the hard palate. Moreover, this feature is detected already in young specimens with a non-overgrown suture between the basisphenoid and the basilar part of the occipital bone (Fig. 9). The shape of the posterior palatine notch demonstrates individual variability. However, in *Spalax* sp. (2n =62), we did not find variants of the posterior palatine notch characteristic of *S. microphthalmus* (“straight edge”) or variants with a small notch in the anterior direction, as in the specimen of *S. zemni* in Fig. 9 or in *S. graecus* (a cleft in the area of the suture between the palatine bones: a trace of the styloid process (Topachevski 1969)). Some specimens of *S. giganteus* also showed weak traces of the styloid process, characteristic of species of the genus *Nannospalax*, but this feature is highly variable in this species.

**Figure 9.**
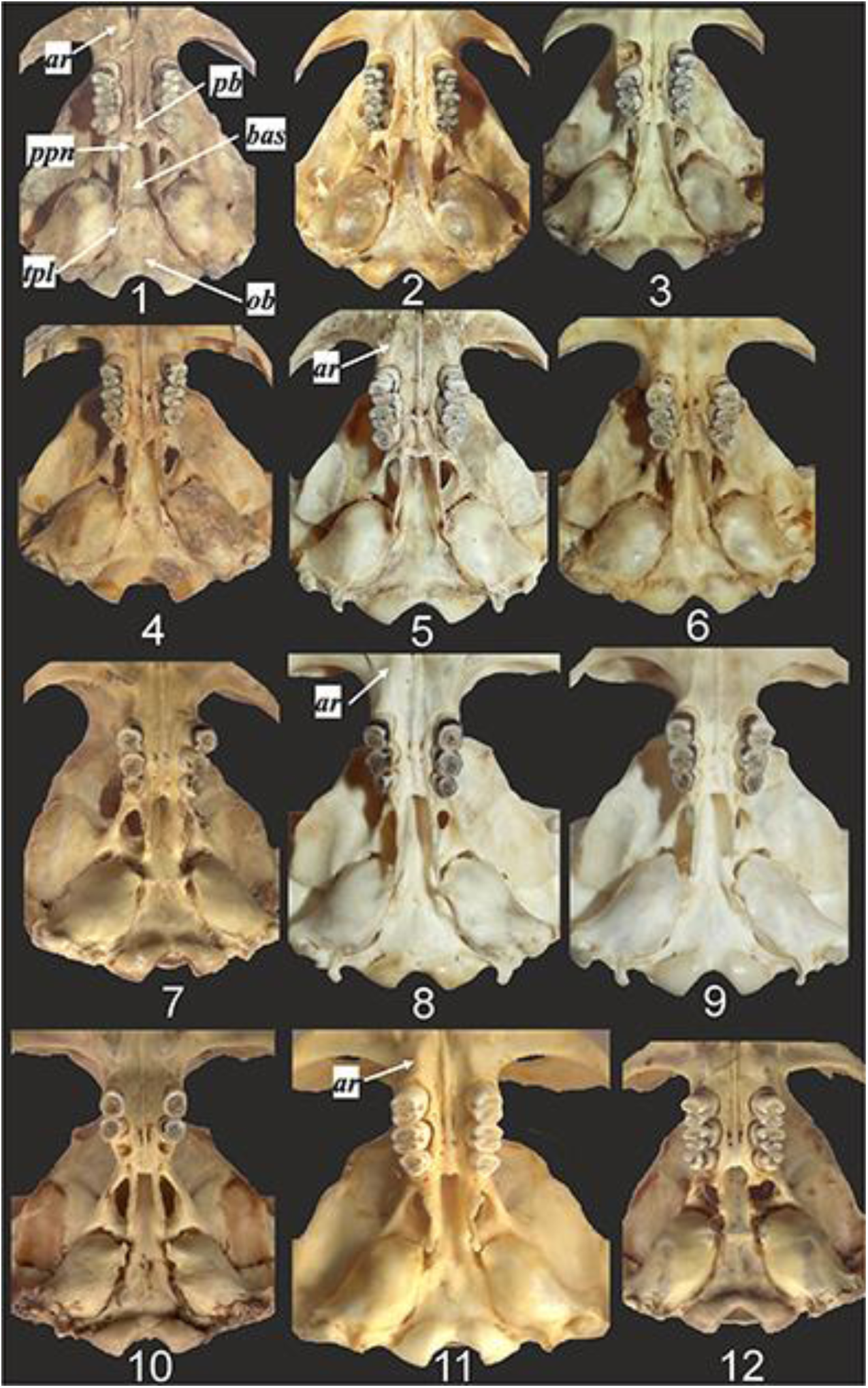
The base of the skull in different species of the genus *Spalax* (drawn to the same scale approximately). 1–6 – *Spalax* sp. (2n = 62): (1) TI S-13610, young female, (2) TI S-10801, young female, (3) KBSU S-3496, young female, (4) IT S-13989, adult male, (5) TI S-1752, adult male, (6) KBSU S-228, adult female; 7 – *S. zemni*, ZIN 66577, adult female, 8, 9 – *S. microphthalmus*, IDB 27272(1), adult male, IDB 27264, adult male, 10 – *S. graecus*, ZIN 66556, adult male, 11 – *S. giganteus*, ZIN 46967, adult female, 12 – *S. uralensis*, ZIN 66584, young female. Abbreviations: *ar* - alveolar ridge, *pb* – palatine bones, *ppn* – posterior palatine notch, *bas* – basisphenoid, *ob* –occipital bone.

The rostral part of the skull in *Spalax* sp. (2n =62) is not widened, in contrast to *S. giganteus*, for example (Fig. 10). The rostrum is wedge-shaped and gradually narrowing towards the front. The average relative width of the rostrum in its middle part does not statistically differ from this measurement in the *S. graecus*, *S. microphthalmus*, and *S. arenarius* (Table S5, Supporting information). The relative length of the nasal bones is similar to that of *S. microphthalmus* and greater than that of the other species. In contrast to *S. microphthalmus*, a notch in the *sutura frontonasalis* is usually present, but it is much smaller than in *S. graecus* (Fig. 10). The posterior edges of the nasal bones are usually slightly “pointed”, in contrast to the blunted edges in *S. microphthalmus* or *S. giganteus*. In adults, the *sutura frontointermaxillaris* is located either at the level of the posterior edge of the nasal bones or slightly behind them, but never as far back as in *S. giganteus*. It is worth noting that in *Spalax* sp. (2n = 62), all the features discussed above are to a greater or lesser extent affected by individual variability.

**Figure 10.**
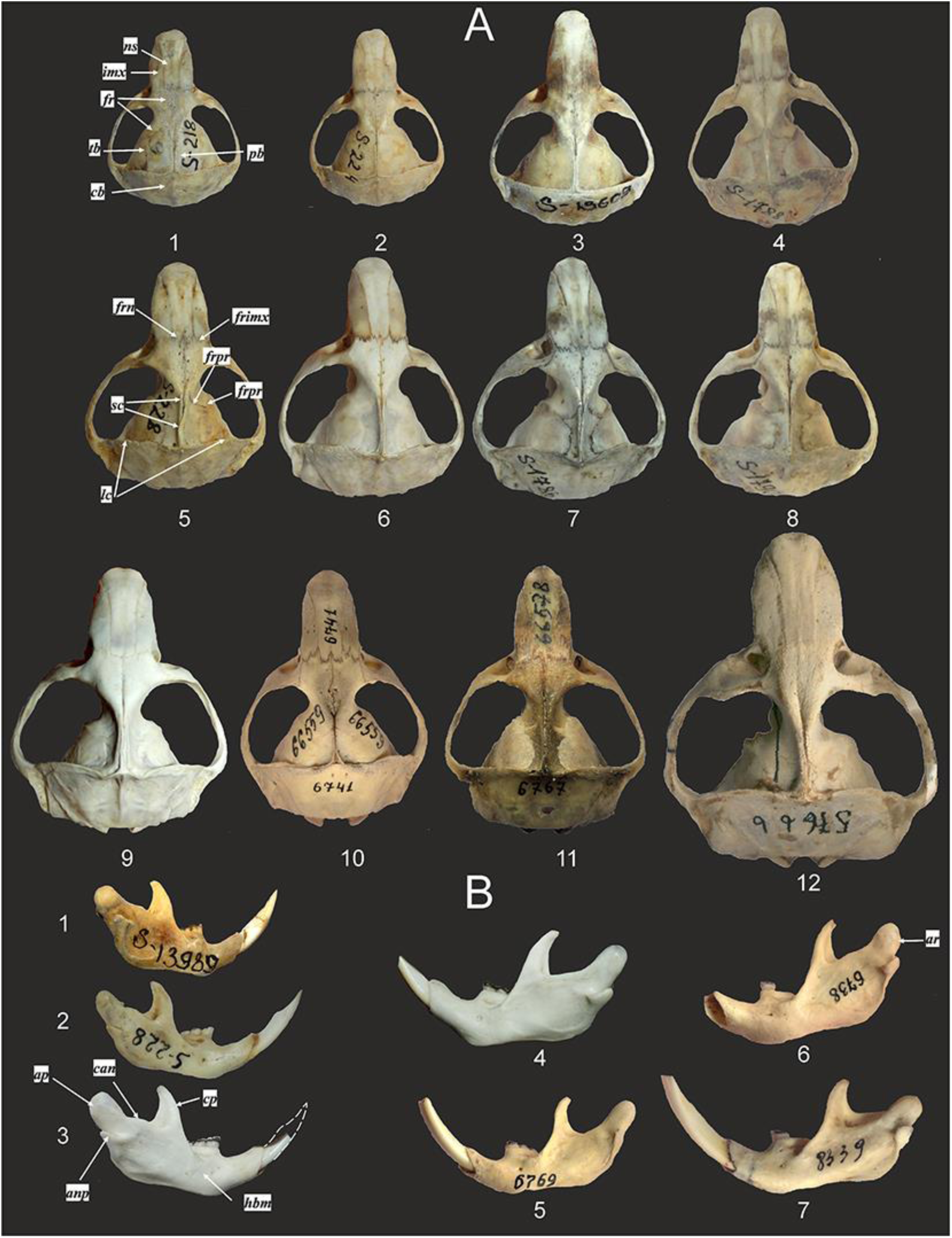
The skulls (A) and mandibles (B) in different species of *Spalax* (drawn to same scale approximately). A: 1–8 *Spalax* sp. (2n = 62) – (1) KBSU S-218, young male, (2) KBSU S-224, young female, (3) TI S-13609, subadult male, (4) KBSU S-1788, adult female, (5) S-228, adult female, (6) IDB 27262, adult male, (7) KBSU S-1786, adult male, (8) KBSU S-1790, adult male; 9 – *S. microphthalmus*, IDB 27271, adult male; 10 – *S. graecus*, ZIN 66559, adult male; 11 – *S. polonicus*, ZIN 66578, adult male; 12 – *S. giganteus*, ZIN 57666, adult female. B: 1–3 *S.* sp. (2n = 62) – (1) TI S-13989, adult female, (2) KBSU S-228, adult female, (3) IDB 27262, adult male; 4 – *S. microphthalmus*, IDB 27272, adult male; 5 – *S. zemni*, ZIN 66577, adult female; 6 – *S. graecus*, ZIN 66556, adult, gender unknown, 7 – *S. uralensis*, ZIN 66580, subadult male. Abbreviations: *ns* – nasal bone, *imx* – intermaxillary bone, *fr* - frontal bone, *pb* – parietal bone, *tb* – temporeal bone, *ob* – occipital bone, *sc* – sagittal crest, *lc* – lambdoidal crest, *frn* – *sutura frontonasalis*, *frimx* – *sutura frontointermaxillaris*, *frpr* – *sutura frontoparietalis*, *frtp* – *sutura frontotemporalis*, *hbn* - horizontal branch of mandible, *ap* – alveolar process, *ar* – alveolar ridge, *cp* – coronoid process, *anp* – angular process, can – corono–alveolar notch.

Age variability of the skull shape and progressive development of the sagittal and lambdoidal crests are typical of the blind mole rats, not only of *Spalax* (Topachevski, 1969). The lambdoidal crest of *Spalax* sp. (2n = 62) is less powerful in comparison with this structure in *S. microphthalmus* and *S. graecus*. The skull height (CPSH) indirectly reflects the degree of development of this crest. The relative skull height in *Spalax* sp. (2n = 62) is about 41.9% of the skull length (38-43.6%). This value corresponds to the relative skull height in *S. giganteus* and *S. uralensis* (41.9% in both species) (Table S5, Supporting information). The other species (excluding *S. zemni*) have a skull height index greater than 43%.

The horizontal branch of the mandible of *Spalax* sp. (2n = 62) was relatively lower (MDH/MNDL = 27.9%, 25.9-31.2%) than in *S. microphthalmus* (29.7%, 23.7-33.5%) and close to *S. graecus* (28.3%, 24.9-32.2%) and *S. arenarius* (27.3%, 23.1-30%). The alveolar process without a ridge on the back surface that presents in *S. graecus* (Fig. 10B: 6) is much higher than the articular process in adult animals. The shape of the corono-alveolar notch (*incisura corono-alveolaris*) is analogous to that of *S. graecus* (Fig. 10B). The irregularity of the notch edge is due to the strong development of the anterior ridge of the alveolar process, which “comes into contact” with the coronal process (Topachevski 1969). Thus, *Spalax* sp. (2n = 62) is the fourth form with this kind of structure of the corono-alveolar notch, together with *S. graecus*, S*. arenarius*, and the extinct *S. minor* (Topachevski, 1969).

There was no difference between *Spalax* sp. (2n = 62) and *S. microphthalmus* in the shape of the upper incisors. However, *Spalax* sp. (2n = 62) had significantly lower INCW/MSL ratios (14.8%) than all the other recent species (Table S5, Supporting information). According to our data (Table 3), the ratio of transverse to longitudinal length of the lower incisor is higher in *S. microphthalmus* than in *Spalax* sp. (2n = 62) (99.5%), but in this case we only consider this as a trend. According to (Topachevski 1969), in *S. giganteus*, the index was 95.2-98.0 (mean)-108.1%. The type series (Nogaisk = Prymorsk, Zaporizhia) of the Early-Middle Pleistocene *S. minor* had INCW/MSL between 90.0-104.1%, mean - 97.7% (N = 20) (Topachevski 1969). The few specimens of *S. minor* from other Pleistocene localities from the lower Dnieper and the Azov-Black Sea region had an index of 88.9-96.6% (Topachevski 1969; Stadnik 2009). In the Middle Pleistocene form *S.* cf. *microphthalmus,* this index is within the range of 93.9-100.5-106.6% (Stadnik 2009), and, therefore, the latter form is closer to the recent *S. microphthalmus* than to *Spalax* sp. (2n = 62). Thus, it is likely that *Spalax* sp. (2n = 62) has the narrowest lower incisors among all the members of *Spalax*, including *S. giganteus* and the extinct *S. minor*.

*Spalax* sp. (2n = 62) has proportionally small molars, which is reflected in the absolute length of the tooth rows, compared with other species of the genus, except for *S. giganteus* and *S. uralensis*. The relative length of the upper and lower tooth rows of *Spalax* sp. (2n = 62) is on average slightly larger than that of *S. microphthalmus* and smaller than that of *S. graecus*. However, no differences are statistically significant (Table S5, Supporting information). The absolute size and shape of the upper molars in *Spalax* sp. (2n = 62) are close to those of *S. microphthalmus* (Table 3; Fig. S8, Supporting information). More differences between these forms have been found in the lower molars. The first and third lower molars in *Spalax* sp. (2n = 62) are wider, and the second lower molar is somewhat narrower than the corresponding molars in *S. microphthalmus*.

The average parameters of the molars in *Spalax* sp. (2n = 62) differ significantly from those typical of *S. minor* (Stadnik 2009) (Fig. 11). However, the minimum values of the length of M1-M3 and m1-m3 and the width of m2 and m3 fall within the range of *S. minor*. The variability in the length of M1 and M2, as well as in the length and width of M3, m1, m2, and m3 in the North Caucasian *Spalax* sp. (2n = 62) covers the variability in the fossil form *S.* cf. *microphthalmus*, dated within a wide interval of ∼0.76-0.37 Ma (Stadnik 2009). At the same time, the latter form exhibited, on average, narrower upper molars, a narrower m1, but a wider m2.

**Figure 11.**
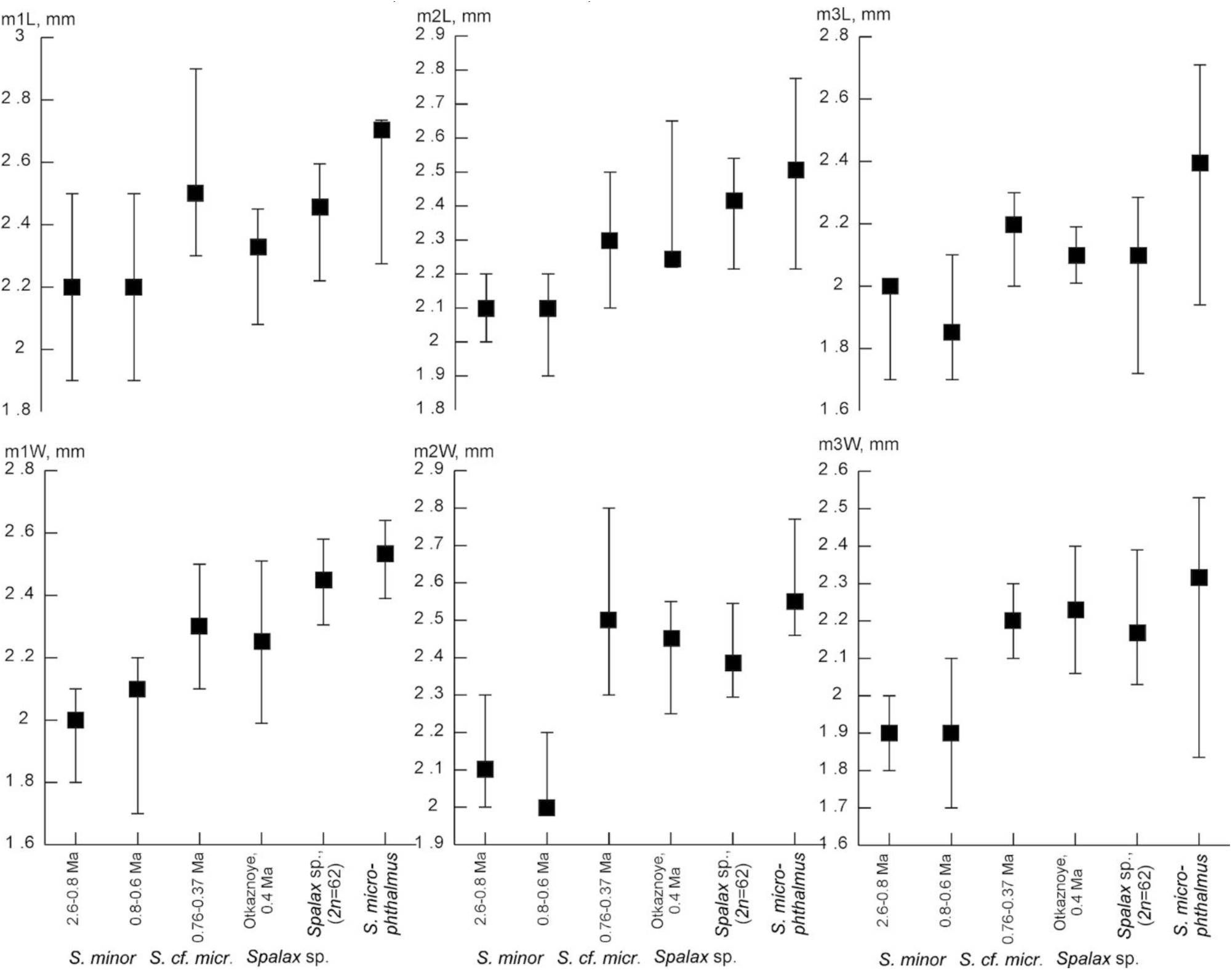
The medians and the minimum-maximum values of the length (L) and width (W) of the upper (M1-M3) and lower (m1-m3) molars of *S. minor* from the Early Pleistocene – the beginning of the Middle Pleistocene from the lower Dnieper and the Azov-Black Sea region, and *S*. cf. *microphthalmus* (Stadnik 2009); *Spalax* sp. from the Otkaznoye locality, the middle of the Middle Pleistocene (material collected and studied by Dr. A.K. Markova); and *Spalax* sp. (2n = 62) (Kabardino-Balkaria, Russia) and *S. microphthalmus* (mainly a population sample from the Kursk district, Russia).

For the lower molars, there is a “trend” of tooth size change from the Early Pleistocene *S. minor* to the modern *S. microphthalmus*. Molars from the Middle Pleistocene sites (*S*. cf. *microphthalmus*, ∼0.76-0.37 Ma) (Stadnik 2009), the Otkaznoe locality, and the recent *Spalax* sp. (2n = 62) occupy an intermediate position (Fig. 11).

The pattern of the molar crown masticatory surface in *Spalax* is very diverse on the one hand and not very different between species on the other. Among all forms of individual and group variability, age-related variability predominates (Topachevski 1969; Puzachenko 1991) (Figs. 3 and 12). The most complex crown structure is observed in juveniles and sub-adult animals, up to 1-1.5 years of age. In these age groups, the structure of crowns clearly exhibits features typical of fossil *Spalax* species such as: separation of the anterocone and paracone on M1, as well as of the proto- and hypocone on all the upper molars and of the proto- and hypoconid on the lower molars; and in addition, there is isolation of the protoconid on m1 (Topachevski 1969; Sarica & Sen 2003). The masticatory surface simplifies greatly with the age of an animal. In addition to age-related variability, the second most important form of molar variability is individual variability. The surface pattern in animals of the same age may not only differ significantly in detail, but may also be different on the left and right tooth rows. Thus, in the small series of molars of young *Spalax* sp. (2n = 62) individuals (N = 12) that was examined, we could not find a clear combination of features that would be completely absent from other extant *Spalax* species or from the Early-Middle Pleistocene *S. minor* (Topachevski 1969; Stadnik 2009). However, there was some discrepancy between the age features of the skull (the development of the sagittal and lambdoidal crests, as well as the shape of parietal bones) and the degree of crown wear: in adult *Spalax* sp. (2n = 62) animals, according to the characteristics of the skull, the crowns were less worn than, for example, in adult *S. microphthalmus* or *S. graecus* specimens.

**Figure 12.**
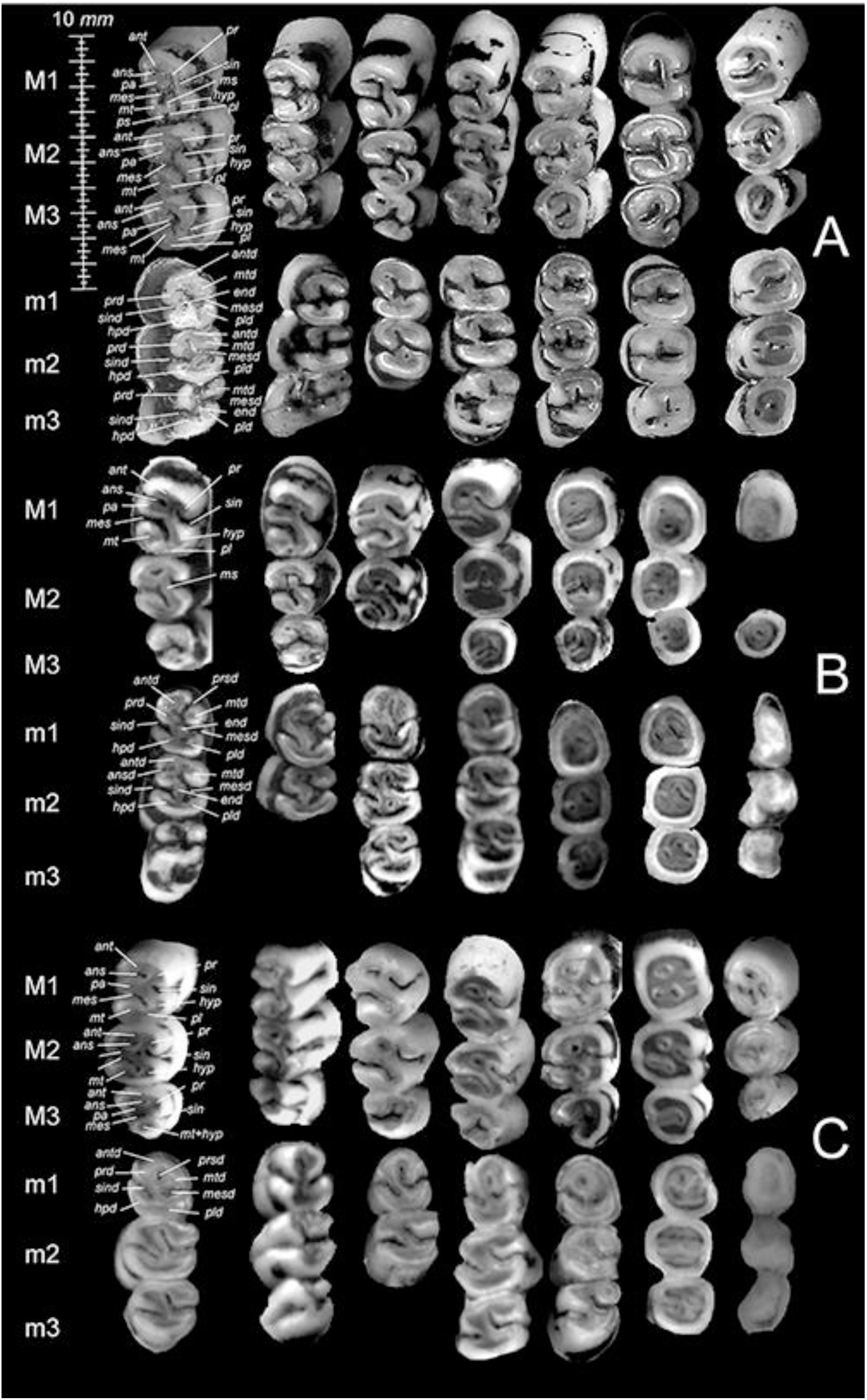
Age-dependent (from left to right) masticatory surface patterns of the upper (M1– M3) and lower (m1–m3) molars in representatives of the genus *Spalax*. A – *Spalax* sp. (2n = 62); specimens from left to write – TI S-13989, S-1752, KBSU S-3495, TI S 1112, S-10801, IDB 27262, and KBSU S-218. B – *S. graecus*; specimens from left to write – ZIN 66561, 65560, 86833, 66555, 66558, 66556, and 66562. C – *S. giganteus*; specimens from left to write – ZIN 180170, n/n, 53621, 53623, 58001, 57666, and 58000. The age variability of the masticatory surface pattern in *S. microphthalmus* and the dental terminology are presented in Fig. 3.

## Discussion

### Karyological features and genetic structure of S. microphthalmus s. l. and the place of the North Caucasian Spalax sp. (2n = 62) in the Spalax phylogeny

The karyotype of North Caucasian blind mole rats (2n = 62) does not differ significantly from other 62-chromosome karyotypes of *Spalax* species. They all have chromosome sets with the same ratio of metacentrics, submetacentrics, and subtelocentrics with very similar morphology (Arslan *et al*. 2016). The sex chromosome morphology of *Spalax* species is also similar, with the exception of *S. giganteus*, in which the X chromosome is unequal-armed (submetacentric or subtelocentric).

*S. microphthalmus* (2n = 60) is the only species with a distinct chromosome set in the karyotypically conservative genus *Spalax*. In the 60-chromosome karyotype, one pair of chromosomes is absent, when compared with the 62-chromosome karyotype. We assumed that these are small submetacentric chromosomes, possibly homologous to the smallest pair of submetacentrics in the *Spalax* sp. set, based on the G-banding data available (Figs. 4B, 5B). It is difficult to determine the location of the translocation of these elements in the 60-chromosome set. This requires the use of high-resolution staining methods. It is quite probable that the 62-chromosome North Caucasian blind mole rat has maintained the karyotype of the ancestral form common to itself and *S. microphthalmus* (2n = 60).

The distribution area of *Spalax* sp. (2n = 62) and its boundary with *S. microphthalmus* (2n = 60) are not clearly defined. Karyotyped individuals with 2n = 62 were found in two localities in Kabardino-Balkaria (Dzuev & Shogenov 2004) and in the south of the Stavropol Krai (our data) in marginal southern populations of *S. microphthalmus* s. l. The nearest localities where blind mole rats with 2n = 60 were found are the right bank of the Manych River and the left bank of the mouth of the

Don River (our data), as well as the Taman Peninsula (Martynova 1977) located at a distance of 330-450 km from the localities of *Spalax* sp. (2n = 62). The question of whether there is a contact zone between 60- and 62-chromosome blind mole rats or whether they are allopatric remains unresolved. It is worth noting that non-karyotyped animals morphologically similar to *Spalax* sp. (2n = 62) and different from *S. microphthalmus* (our data) were previously captured in the territory of Kabardino-Balkaria from the foothills to an altitude of about 2000 m a.s.l. (Dzuev and Shogenov 2004; Tembotova 2015).

The position of *Spalax* sp. (2n = 62) on the phylogenetic tree (Fig. 6) and the high values of genetic distances (Table 2) differentiating it from all the other *Spalax* species, including *S. microphthalmus* and *S. giganteus* with which it was previously associated (Dzuev & Shogenov 2003; Korobchenko and Zagorodniuk 2009), indicate that this form represents a separate phyletic lineage of blind mole rats with an independent evolutionary history. The branch lengths in the clade of the sister species *Spalax* sp. (2n = 62) and *S. microphthalmus* and the topology of the haplotype network (Fig. 7) testify to their origin from a common ancestral form, and at the same time, neither of them is parental to the other. Based on the combination of genetic features studied, we can generally conclude that *Spalax* sp. (2n = 62) has achieved species-level genetic differentiation from the other *Spalax* species.

The branching of the *Spalax* tree indicates the initial division of large-bodied blind mole rats into two phyletic groups. One of these groups includes the western *S. antiquus* and *S. graecus*, while the other includes all the other species distributed to the east. Given the divergence dates obtained by Hadid *et al*. (2012), according to which the *antiquus*/*graecus* divergence time is ∼1.3 Ma, the *xanthodon*/*leucodon* divergence time is ∼2.7 Ma, and the *Spalax*/*Nannospalax* divergence time is ∼7.6 Ma, the divergence time of *Spalax* sp. (2n = 62) and *S. microphthalmus* (2n = 60) can be extrapolated as ∼1.7 Ma, based on the tree topology. A plausible paleogeographic event that may have contributed to the isolation of these two forms of blind mole rats could be the Apsheron Ponto-Caspian transgression, which separated the Caucasus from the Eastern European Plain for a period of approximately 500,000 years (1.8-1.3 Ma) (Forte & Cowgill 2013; Svitoch 2016).

The low intraspecific genetic variability of *S. microphthalmus* (2n = 60) and the absence of an explicit phylogenetic structure may indicate the low efficiency of potential ecological and geographical barriers in the territory of its distribution. Another alternative may be the recent rapid colonization of the area by a local ancestral population. The combination of a high value of haplotype diversity (*Hd*) with a low value of nucleotide diversity (*π*) suggests that the recent population may have originated from an ancestral population with a low effective population size. A rapid expansion of the population is also indicated by the relatively high value of the expansion coefficient (S/k). The above data, combined with a low level of intraspecific genetic variability and the absence of geographic subdivision, suggest that the modern range of *S. microphthalmus* was formed as a result of rapid dispersal from one or several small refugia in historically recent times. This process may have begun during the Late Pleistocene, coinciding with the formation of the periglacial steppes of Europe, and may have subsequently intensified during the Holocene. The abundance of Holocene fossils of *S. microphthalmus* (Topachevski 1969) may serve as indirect evidence of this expansion.

### Morphometric and morphological features of the North Caucasian Spalax sp. (2n = 62) in the genus Spalax

The results of morphometric analysis and comparison of modern and fossil representatives of the genus *Spalax* allowed us to reveal a unique combination of qualitative and quantitative features in *Spalax* sp., distributed in the central part of the North Caucasus. The sample used for morphometric and morphological studies included skulls of animals captured in the region of interest (Kabardino-Balkaria) and skulls stored in collections. A preliminary comparison of the latter with the skulls of karyotyped specimens showed their similarity in characteristic features (the structure of the hard palate, the small size of adult specimens, etc.). Further analysis confirmed the relative compactness of this geographic sample.

In the skull structure of the North Caucasian *Spalax* sp., some features stand out that may reflect an adaptation to digging, compared with *S. micropthtalmus*, *S. arenarius*, etc. A rough estimate of the angle between the left/right *fossa glenoidea* (Fig. S8A, Supporting information) is approximately 18.8°, while in other modern species this value exceeds 19° (19.7° to 23.8°). According to G.E Zubtsova (Zubtsova 1986), before blind mole rats bite the soil, the condyle of their condyloid process is within the middle third of the *fossa glenoidea*; and in this position of the mandible, the lower incisors are spread apart (Fig. S7B, Supporting information). The biological interpretation could be as follows: a) *Spalax* sp. (2n = 62) is more adapted to digging in dense soils; or b) this species generally has a relatively low adaptation to digging among all the other *Spalax* species. We cannot rule out compensation for the described feature of the skull base with a relatively long condylar process, which slightly increases the overall length of the lower jaw. In addition, this form has a relatively narrow articular head (mean of length/width ratio: 28.5%), in comparison with *S. giganteus* (38%), *S. microphthalmus* (31.1%), and *S. graecus* (30.1%), which should slightly limit the possibility of lateral mandible movements. This limitation, together with the sharper angle between the glenoid cavities, suggests a relatively small divergence of the upper parts of the lower incisors during digging, which reduces their overall efficiency (Zubtsova 1986).

The degree of development of the lambdoid crest is positively related to the development of the *musculus rhomboideus capitis*, which is attached along the entire length of this crest and terminates at the scapula (Gambaryan 1960). The strength of this muscle is related to the magnitude of the forces generated when the animal raises its head, which is of great importance in the digging process. The relatively low lambdoid crest in *Spalax* sp. (2n = 62) indicates a lower (for *Spalax*) specialization of the skull for digging activities. The development of the crest is proportional to the load at the site of muscle attachment, and the crest itself increases with age, reaching its maximum size (in all species) in adult and old males. It should be noted that this process seems to take longer in ontogeny in *Spalax* sp. (2n = 62) than in *S*. *microphthalmus*. For this reason, sexually mature specimens of *Spalax* sp. (2n = 62) caught in Kabardino-Balkaria were often previously classified as subadults of *S. microphthalmus*. The relatively narrow incisors of *Spalax* sp. (2n = 62), compared with all the other modern species, may also indicate a low digging efficiency. However, the same features can also be considered in the context of an adaptation to dense soils containing stone fragments or debris, which is typical of the known range of *Spalax* sp. (2n = 62).

The probable origin of *Spalax* sp. (2n = 62) can be traced back at least to the middle of the Middle Pleistocene of Eastern Europe. Considering the geographical position of the Otkaznoe locality, we assumed that the *Spalax* sp. found there is the direct ancestor of the modern *Spalax* sp. (2n = 62) in the region. The available data do not exclude the possibility of a wider distribution of this form in Eastern Europe during the Middle Pleistocene. In our opinion, a revision of the fossil remains of “*S*. cf. *microphthalmus*” from the Middle-Late Pleistocene deposits is necessary.

In general, the combination of original morphological features and small size (the smallest among the modern *Spalax* species) allows us to assume that the North Caucasian blind mole rat is a distinct species within the genus *Spalax*. This assumption is supported by the available genetic data.

#### Taxonomic section

The combined data presented above suggest that the North Caucasian blind mole rat with karyotype 2n = 62, previously classified as the greater blind mole rat, *S. microphthalmus* (Dzuev 1989; Dzuev & Shogenov 2003; Dzuev *et al*. 2019) or *S.* cf. *giganteus* (Korobchenko & Zagorodniuk 2009), is a new species of the genus *Spalax*.

Class: Mammalia Linnaeus, 1758

Order: Rodentia Bowdich, 1821

Suborder: Myomorpha Brandt, 1855

Superfamily: Muroidea Illiger, 1811

Family: Spalacidae Gray, 1821

Subfamily: Spalacinae Gray, 1821

Genus: *Spalax* Güldenstädt, 1770

***Spalax lyapunovae*** sp. nov. Brandler, Tukhbatullin, Kapustina, Puzachenko

*Spalax microphthalmus.* Tembotova, 2015: Fig. 129

*Spalax microphthalmus*. Tembotov and Shkhashashev, 1987: 172-173

*Spalax microphthalmus*. Dzuev, 1989: 55-57

*Spalax microphthalmus*. Dzuev *et al*., 2025a.

*Spalax* cf. *giganteus, Spalax* ex gr. *Giganteus-arenarius*, «giganteus» group. Korobchenko and Zagorodniuk, 2009: 16, 22

#### Local name

Lyapunova’s blind mole rat.

#### Holotype

S-211554, an adult/old male specimen preserved as skull and skin (Fig. 13) in good condition is held in the Zoological Museum of Lomonosov Moscow State University (ZMMU) (Moscow, Russia), and tissue samples are stored in the ‘Collection of wildlife tissues for genetic research’ of the Koltzov Institute of Developmental Biology of the Russian Academy of Sciences (CWT IDB) (Moscow, Russia) under collection ID 27260; collected by O.V. Brandler, A.R. Tukhbatullin, and S.Y. Kapustina on 16 May 2021. The GenBank accession number PV012729 provides the mitochondrial cytochrome *b* sequence.

**Figure 13.**
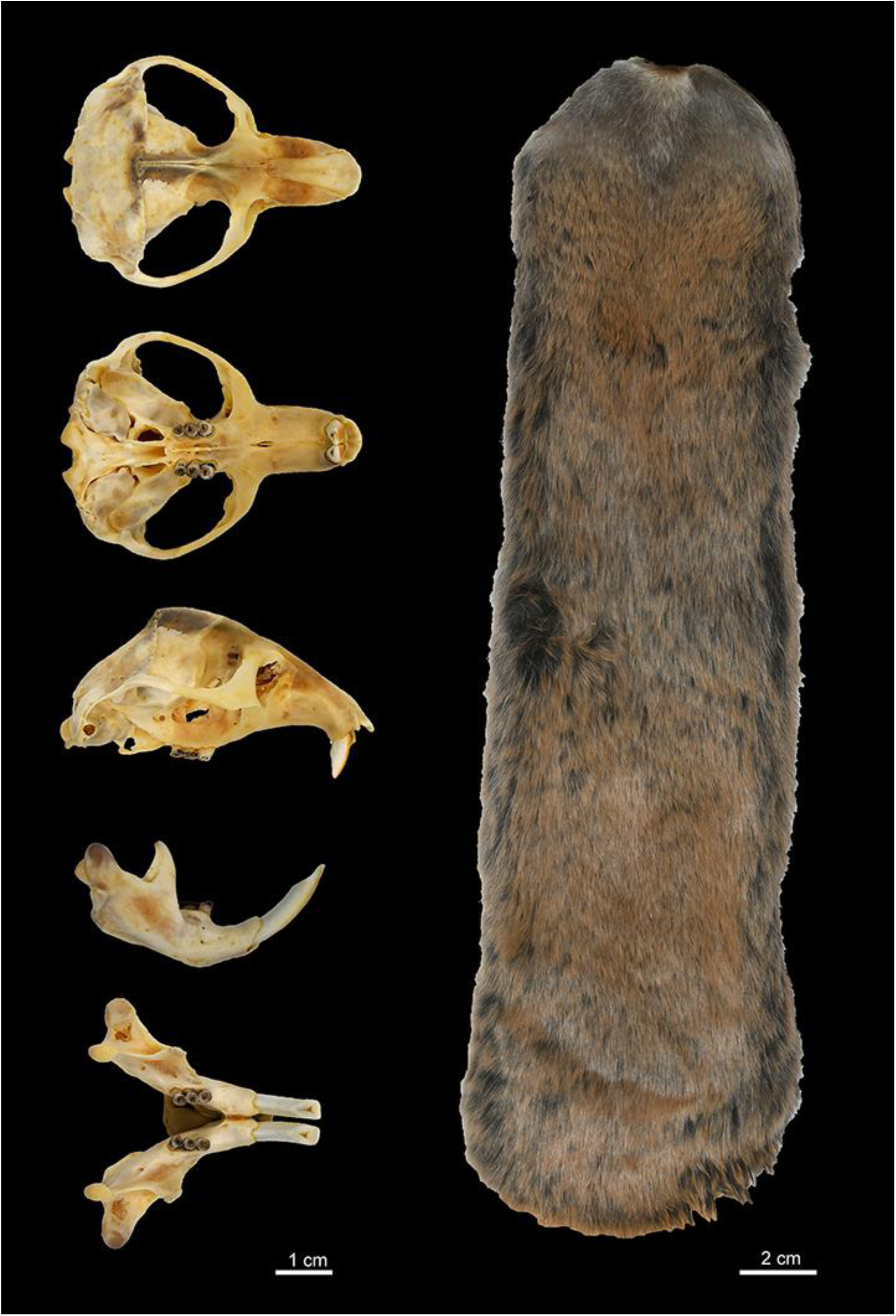
The skull and skin of the *Spalax lyapunovae* sp. nov. holotype. Museum ID: S-211554 (ZMMU, Moscow, Russia).

#### Type locality

7 km south-south-west of Kislovodsk (43.835324°N, 42.682150°E, elevation ∼ 1382 m above sea level (a.s.l.), coordinates taken by GPS), Predgorniy Municipality, Stavropol Krai, Russia.

#### Paratypes

Five specimens (skulls and skins in ZMMU and tissue samples in CWT IDB) collected by O.V. Brandler, A.R. Tukhbatullin, and S.Y. Kapustina in the type locality: 1) S-211553 ZMMU, 27259 CWT IDB, adult male, 15 May 2021; 2) S-211555 ZMMU, 27261 CWT IDB, adult female, 16 May 2021; 3) S-211556 ZMMU, 27262 CWT IDB, adult male, 16 May 2021; 4) S-211557 ZMMU, 27263 CWT IDB, subadult male, 16 May 2021; and 5) S-211558 CWT 27268 IDB, adult male, 16 May 2021. The GenBank accession number PV012730 provides the *cytb* sequence common to all paratypes. Seven specimens (skulls and skins) are housed in the Kabardino-Balkarian State University (KBSU) (Nalchik, Russia) and Tembotov Institute (TI) (Nalchik, Russia) collections: KBSU S-1786 and KBSU S-1788, adult male and adult female respectively, skulls, “Aursentkh” (“Narzan Valley”) (approximately 43.695582°N, 42.677053°E, elevation ∼2000 m a.s.l.), Zolsky District, Kabardino-Balkaria, Russia, collected by G.I. Dzuev and A.L. Shogenova, 3.08.2003; KBSU S-1790, adult male, skull, from the same place, collected by G.I. Dzuev and A.L. Shogenova, 5.08.2003; KBSU S-228, adult female, skull, 6 km west of Prokhladny city (dacha areas) (approximately 43.773479°N, 43.905801°E, elevation ∼150 m a.s.l.), Prokhladny city district, Kabardino-Balkaria, Russia, collected by A. Lenshin, 25.06.99; TI S-13609, adult male, skull, Psynadakha village (approximately 43.843078°N, 43.218281°E, elevation ∼650 m a.s.l.), Zolsky district, Kabardino-Balkaria, Russia, collected by A.A. Tembotov and F.A. Tembotova, 15.06.2000; TI S-13382, juvenile female, skull, the same place of trapping, collected by F.A. Tembotova, 15.06.2000; TI S-10801, “Aursentkh” (approximately 43.695582°N, 42.677053°E, elevation according to the museum label: 2000 m a.s.l.), Zolsky District, Kabardino-Balkaria, Russia, collected by G.I. Dzuev, 25.07.1987.

#### Diagnosis

A typical member of the genus *Spalax* as diagnosed by V.A. Topachevski (Topachevski 1969). *S. lyapunovae* may be recognized by the following combination of characteristics. It is the smallest living species in the genus. The skull is relatively low with a wide base. The rostrum is relatively high and narrow in the posterior part. The *foramen incisivum* is very short. The interorbital width of the skull is relatively large. The posterior palatal notch is convex, more or less rounded. The *fossa glenoidea* is extraordinarily short. The mandible is massive and relatively long; the *incisura corono-alveolaris* has an irregular margin. The upper and lower incisors are narrow.

#### Karyotype

Diploid chromosome number 2n = 62 (*NFa* = 120, *NF* = 124).

#### Description

Body weight (all ages): 221.4±16.6 g (120 – 315 g, *N* = 15), hind foot length: 24.8±0.3 mm (22.1 – 26.3 mm, *N* = 15). According to R.I. Dzuev *et al*. (Dzuev *et al*. 2019; 2025b), blind mole rats from the northern slope of the Central Caucasus have a body mass ranging from 170 g to 350 g, a body length ranging from 167.0 to 239.5 mm, and a hind foot length ranging from 19.0 to 27.2 mm.

The *S*. *lyapunovae* sp. nov. has an apparently unique convex shape of the posterior edge of the palate (palatine notch). The maximum skull length (MSL, adult animals) constitutes 50.4±0.47 mm (47.2–5.2 mm, *N* = 18). The mastoid width (MSW) is 52.2±0.32% (50.3–55.1%) of the maximum skull length and is on average larger than in other species of the genus. The posterior rostrum height (RSMAH) is 77.4±0.47 % (67.2–88.2 %) of the rostrum length (RSL) and 25.6±0.35 % (22.7–27.9 %) of the maximum skull length, which is higher than in all the other species of the genus, except for the Ural blind mole rat (*S. uralensis*). The skull width between the orbits (YW) in relation to the skull length (15.4±0.23 %, 13.1-17.2%) is on average greater than in other *Spalax* species. The species has probably the smallest alveolar length of the upper tooth rows (7.67±0.09 mm, 6.97–8.56 mm), but especially of the lower tooth rows (7.07±0.07 mm, 6.72-7.83 mm), among all recent *Spalax* species. The length of the incisive foramina is very short (3.48±0.10 mm, 3.1–3.8 mm). The average length of the *foramen incisivum* in relation to the length of the upper diastema is 18.7 %. The absolute length of the *fossa glenoidea* (9.27±0.2 mm, 8.11-10.9 mm) and its length in relation to the skull length (18.4±0.29 %, 16.1-20.8%) are the smallest among all the species of the genus. On average, the ratio of transverse and longitudinal sections of an incisor at the level of its alveoli (99.5±1.91 %, 92.9-104.4%, N = 6) is lower than in most members of the genus, with the exception of the giant blind mole rat *(S. giganteus*) and the Early Pleistocene Nogai blind mole rat (*S. minor*).

The chromosome set is characteristic of the vast majority of species belonging to the genus *Spalax*. It is comprised of five pairs of medium and small metacentrics, 12 pairs of submetacentrics decreasing in size, and 13 pairs of subtelocentrics, including two pairs of the largest elements of the set. The X chromosome is a large metacentric or submetacentric, and the Y chromosome is a small subtelocentric.

The complete *cytb* sequence is distinct from those of the other *Spalax* species due to the following unique substitutions: C>A in locus 28, A-G in 294, C>T in 324, C/A>T in 531, A/G/T>C in 462, A/C>T in 126, A/C/T>G in 396, A/C>T in 531, A>G in 888, G>A in 904, and A>G in 990.

#### Comparisons and comments

A comparison with other recent and extinct taxa belonging to *Spalax* is given above. The new species shares several cranial features with *S. graecus*, *S. microphthalmus*, *S. giganteus*, and the extinct Early-Middle Pleistocene *S. minor* and *S*. cf. *microphthalmus.* The species thus occupies a distinctive position in terms of its cranial characteristics. Taking into account the phylogenetic interpretation, the new species is characterized by a set of “primitive” features proposed for a common ancestor of the “western” and “eastern” lineages within the genus *Spalax*. We hypothesize that the new species shares a common ancestor with the greater blind mole rat (*S. microphthalmus*), but is not the ancestor of the latter, and vice versa. The species probably evolved no later than the mid-Middle Pleistocene, before the 60-chromosome form (*S*. *microphthalmus*) evolved from the hypothetical 62-chromosome *Spalax* in Eastern Europe.

#### Etymology

The species is named in honor of Dr. Elena A. Lyapunova, in recognition of her contribution to the study of the genetics and cytogenetics of Spalacidae.

#### Distribution and ecology

The range limits of the new species need to be clarified. It is not known whether the range of the new species overlaps with that of the greater blind mole rat. So far, it can be stated that the species inhabits the Central Caucasus from the foothills (150 m a.s.l) to the subalpine belt (>2000 m a.s.l) (Fig. 1). According to R.I. Dzuev *et al*. (Dzuev *et al*. 2019), the blind mole rats have entered the subalpine belt relatively recently. The change in habitat conditions has affected a number of dimensional and physiological parameters of the animals (body weight, blood parameters, etc.). The ecology of the species itself remains largely unexplored. There is little or no information about the density of its local populations (Dzuev *et al*. 2025a).

## Conclusion

With the recognition of *S. lyapunovae* sp. nov. as a distinct species, the number of extant species of the genus *Spalax* has increased to nine. In addition to the new species, it includes one 60-chromosomal *S. microphthalmus* and seven 62-chromosomal species: *S. giganteus*, *S. uralensis*, *S. zemni* (= *S. polonicus*), *S. arenarius*, *S. graecus*, and after Németh *et al*. (2013), *S. antiquus* Méhely, 1909 and *S. istricus* Méhely, 1909.

A unique combination of morphological characteristics has been described for the new species, including features that bring it closer to the hypothetical ancestor of the modern *Spalax* species. It is necessary to study the range and the ecology of the new species, which is endemic to the central part of the North Caucasus.

### Nomenclatural acts registration

The electronic version of this article in portable document format represents a published work according to the International Commission on Zoological Nomenclature (ICZN), and hence the new names contained in the electronic version are effectively published under that Code from the electronic edition alone (see Articles 8.5–8.6 of the Code). This published work and the nomenclatural acts it contains have been registered in ZooBank (https://zoobank.org/), the online registration system for the ICZN. The ZooBank Life Science Identifier (LSID) for this publication is urn:lsid:zoobank.org:pub:92A576C0-7E24-4415-ACB2-DF2D066818A9.

## Acknowledgments

We would like to thank E.P. Kononenko (TI), A.Yu. Paritov, and M.A. Khashkulova (KBSU), I.Y. Pavlinov and E.L. Yakhontov (ZMMU), and G.I. Baranova (ZIN) for assistance in accessing the collection materials. We would like to extend special thanks to A.K. Markova for so kindly providing the material from the Otkaznoye locality. We thank V.V. Stakheev (Southern Scientific Centre) for his help in collecting in the field. We thank A.V. Andreychev (Mordovia State University) for providing the specimens from Mordovia. We thank Dr. Natasha Grigorian for her help with proofreading the manuscript. The research was conducted as part of the Institute of Geography of the Russian Academy of Sciences State Assignment, Project No. FMWS-2024-0007 (AP) and was funded partly by the Government basic research program No. 0088-2024-0011 of the Koltzov Institute of Developmental Biology, Russian Academy of Sciences (OB, AT, SK).

## Notes

### Competing Interest Statement

The authors have declared no competing interest.

