## Supplementary Material 2 for "Genetical and morphological variability of the greater blind mole rat *Spalax microphthalmus* and phylogenetic affinities of large-bodied spalacids (Rodentia, Spalacidae) with a description of *Spalax lyapunovae* sp. nov. from the North Caucasus"

**Supplementary materials for the section *Cytb variability in the genus Spalax*** ........................ 2

**Supplementary materials for the section *Craniometrical variability in the genus Spalax*** ....... 3

**Supplementary materials for the section *A morphological comparison of the North Caucasian Spalax sp. (2n = 62) with members of the genus Spalax*** ............................................................ 6

**Supplementary materials for the section *Cytb variability in the genus Spalax***

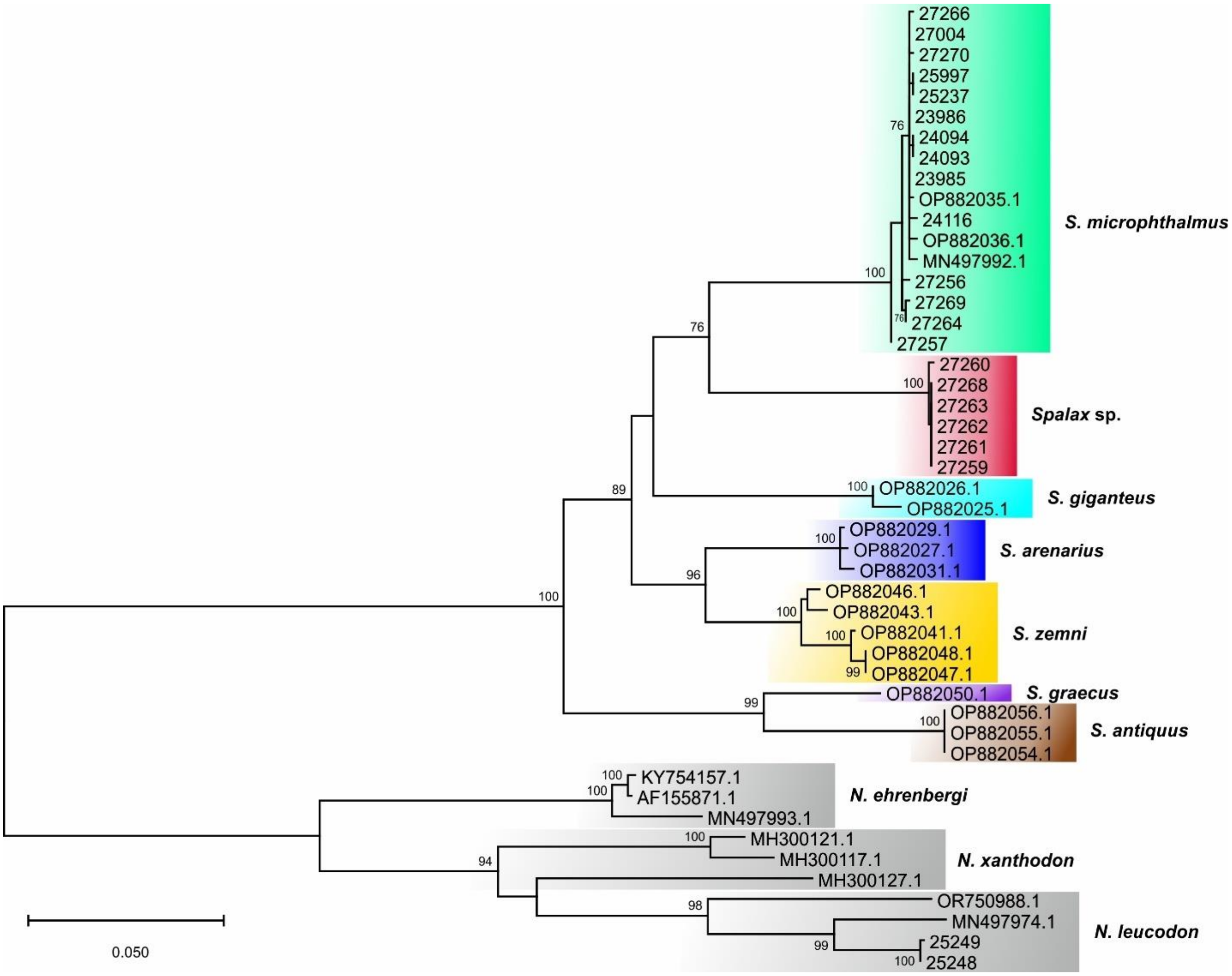


**Figure S1** Phylogenetic ML tree of blind mole rats based on the mitochondrial *cytb* gene. The values at the nodes represent bootstrap support. Values below 70% (ML) and for minor nodes are not shown. The colors correspond to Fig. 6B.

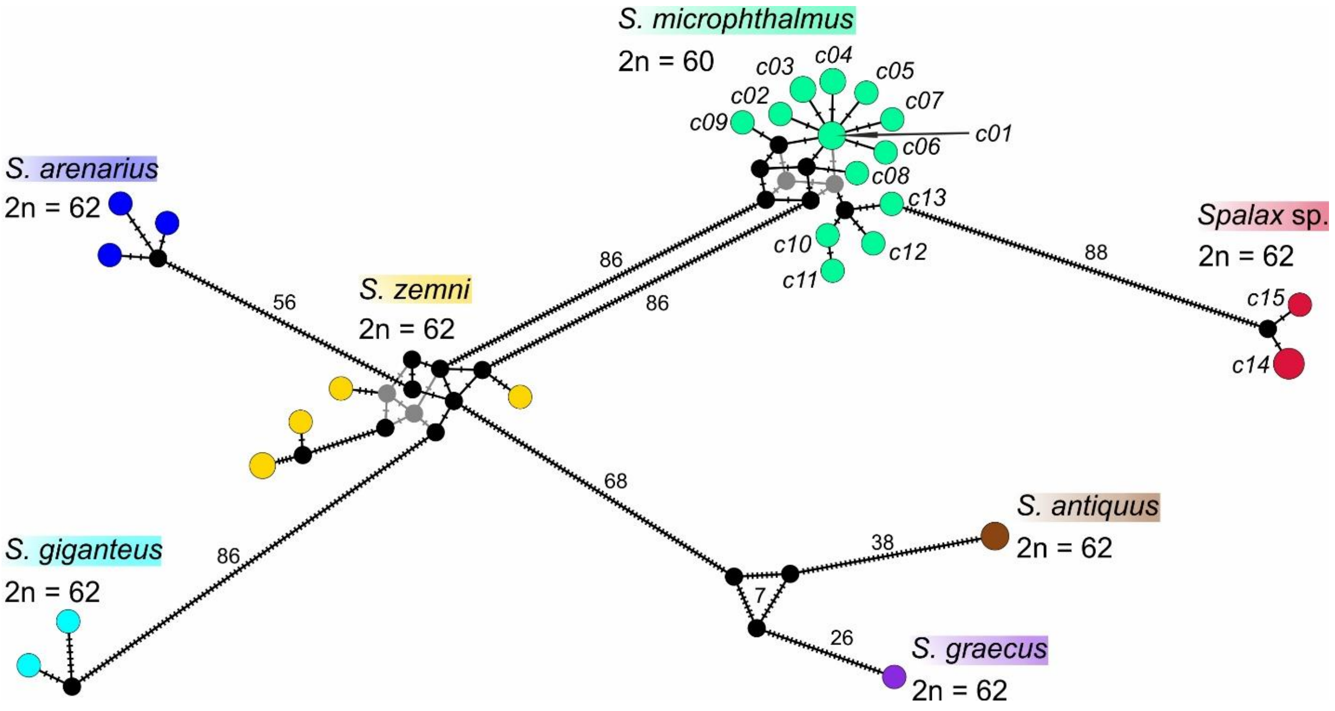


**Figure S2** Evolutionary *cytb* haplotype networks of *Spalax*. Haplotype labels of *S. microphthalmus* and *Spalax* sp. correspond to Table S1, Supporting information. The colors correspond to Fig. 6B.

### Supplementary materials for the section *Craniometrical variability in the genus Spalax*

**Table S3** Pearson correlations/partial correlations of morphometric variables (VAR) with SZM coordinates E1–E4 and indices VAR/ML (except ML) with SHM coordinates K1–K3 in the genus *Spalax* as well as relative components of variance due to putative taxonomy (TAX) or sexual dimorphism (SD); statistically significant *r* values ($p < 0.001$) are highlighted in red

| VAR | SZM model | | | | | VAR/ | SHM model | | | |
|---|---|---|---|---|---|---|---|---|---|---|
| | E1 | E2 | E3 | E4 | $r^2$ | MSL | K1 | K2 | K3 | $r^2$ |
| MSL | 0.98 | -0.05 | -0.01 | 0.09 | 0.97 | ML | 0.56 | 0.46 | -0.46 | 0.64 |
| | 0.98 | -0.29 | -0.02 | 0.30 | | | 0.64 | 0.53 | -0.57 | |
| RSMIH | 0.98 | -0.04 | -0.02 | 0.03 | 0.95 | RSMIH | 0.57 | 0.54 | -0.24 | 0.59 |
| | 0.98 | -0.22 | -0.06 | -0.02 | | | 0.63 | 0.59 | -0.28 | |
| MSW | 0.97 | 0.03 | 0.01 | 0.02 | 0.94 | MSW | 0.15 | 0.09 | 0.38 | 0.18 |
| | 0.97 | 0.08 | 0.09 | -0.03 | | | 0.17 | 0.12 | 0.40 | |
| ZW | 0.97 | -0.08 | 0.02 | 0.07 | 0.96 | ZW | 0.25 | 0.50 | -0.27 | 0.34 |
| | 0.98 | -0.38 | 0.12 | 0.20 | | | 0.23 | 0.49 | -0.27 | |
| RSMEW | 0.97 | 0.11 | -0.11 | 0.02 | 0.96 | RSMEW | 0.77 | 0.32 | -0.14 | 0.66 |
| | 0.98 | 0.44 | -0.44 | -0.02 | | | 0.79 | 0.38 | -0.17 | |
| RSMAW | 0.97 | 0.04 | -0.04 | -0.02 | 0.94 | RSMAW | 0.61 | 0.41 | -0.01 | 0.49 |
| | 0.97 | 0.13 | -0.11 | -0.18 | | | 0.63 | 0.44 | 0.06 | |
| PL | 0.97 | -0.08 | 0.01 | 0.10 | 0.95 | PL | -0.16 | 0.08 | -0.13 | 0.00 |
| | 0.97 | -0.34 | 0.08 | 0.29 | | | -0.17 | 0.08 | -0.13 | |
| FGMEW | 0.96 | 0.15 | -0.05 | 0.05 | 0.94 | FGMEW | 0.39 | -0.19 | 0.34 | 0.31 |
| | 0.97 | 0.51 | -0.14 | 0.10 | | | 0.45 | -0.24 | 0.37 | |
| INCW | 0.95 | 0.11 | -0.19 | 0.05 | 0.95 | INCW | 0.79 | 0.26 | -0.30 | 0.72 |
| | 0.98 | 0.42 | -0.65 | 0.13 | | | 0.82 | 0.29 | -0.44 | |
| FGMAW | 0.95 | 0.17 | -0.04 | 0.09 | 0.94 | FGMAW | 0.42 | -0.27 | 0.30 | 0.36 |
| | 0.97 | 0.57 | -0.14 | 0.26 | | | 0.49 | -0.34 | 0.35 | |

| VAR | SZM | model | | | | VAR/ | SHM | model | | |
|---|---|---|---|---|---|---|---|---|---|---|
| | E1 | E2 | E3 | E4 | $r^2$ | MSL | K1 | K2 | K3 | $r^2$ |
| MNDL | 0.95 | -0.15 | 0.07 | -0.02 | 0.93 | MNDL | -0.23 | 0.33 | 0.29 | 0.27 |
| | 0.97 | -0.52 | 0.28 | -0.19 | | | -0.29 | 0.40 | 0.34 | |
| DUL | 0.94 | -0.22 | 0.03 | 0.15 | 0.95 | DUL | -0.33 | 0.23 | -0.49 | 0.42 |
| | 0.97 | -0.70 | 0.12 | 0.44 | | | -0.43 | 0.29 | -0.54 | |
| FGMIW | 0.94 | 0.13 | 0.00 | -0.02 | 0.89 | FGMIW | 0.24 | 0.03 | 0.40 | 0.23 |
| | 0.94 | 0.35 | 0.02 | -0.14 | | | 0.28 | 0.04 | 0.42 | |
| FCL | 0.92 | -0.17 | 0.07 | 0.19 | 0.91 | FCL | -0.28 | 0.01 | -0.25 | 0.14 |
| | 0.95 | -0.50 | 0.22 | 0.47 | | | -0.29 | 0.02 | -0.27 | |
| RSL | 0.91 | 0.00 | 0.02 | 0.26 | 0.87 | RSL | 0.10 | -0.20 | -0.09 | 0.00 |
| | 0.93 | 0.00 | 0.07 | 0.54 | | | 0.12 | -0.22 | -0.10 | |
| CPSH | 0.91 | -0.15 | 0.06 | 0.09 | 0.86 | CPSH | -0.30 | -0.02 | -0.04 | 0.09 |
| | 0.93 | -0.38 | 0.17 | 0.15 | | | -0.30 | 0.01 | -0.05 | |
| MNDH | 0.91 | -0.16 | 0.10 | -0.07 | 0.88 | MNDH | 0.02 | 0.60 | -0.03 | 0.36 |
| | 0.93 | -0.44 | 0.30 | -0.29 | | | -0.05 | 0.60 | 0.02 | |
| CNDL | 0.91 | 0.01 | -0.02 | 0.12 | 0.84 | CNDL | 0.19 | -0.03 | 0.01 | 0.00 |
| | 0.91 | 0.00 | -0.02 | 0.22 | | | 0.19 | -0.05 | 0.01 | |
| RSMAH | 0.90 | -0.11 | 0.08 | -0.10 | 0.84 | RSMAH | -0.01 | 0.32 | 0.27 | 0.19 |
| | 0.91 | -0.12 | 0.21 | -0.31 | | | -0.04 | 0.36 | 0.31 | |
| RSMIW | 0.89 | -0.10 | 0.12 | -0.19 | 0.87 | RSMIW | 0.11 | 0.61 | 0.18 | 0.42 |
| | 0.93 | -0.30 | 0.35 | -0.05 | | | 0.07 | 0.63 | 0.29 | |
| RSMEH | 0.89 | -0.08 | 0.08 | 0.15 | 0.83 | RSMEH | 0.08 | 0.11 | -0.17 | 0.00 |
| | 0.91 | -0.19 | 0.20 | 0.27 | | | 0.07 | 0.09 | -0.16 | |
| TUL | 0.89 | 0.31 | -0.21 | -0.10 | 0.93 | TUL | 0.80 | 0.06 | 0.20 | 0.69 |
| | 0.96 | 0.75 | -0.60 | -0.42 | | | 0.82 | 0.00 | 0.38 | |
| FGL | 0.88 | 0.10 | -0.15 | 0.24 | 0.84 | FGL | 0.36 | -0.34 | -0.33 | 0.39 |
| | 0.91 | 0.23 | -0.34 | 0.48 | | | 0.45 | -0.46 | -0.41 | |
| TDL | 0.88 | 0.34 | -0.19 | -0.11 | 0.93 | TDL | 0.73 | -0.01 | 0.35 | 0.67 |
| | 0.96 | 0.78 | -0.56 | -0.44 | | | 0.79 | -0.10 | 0.55 | |
| CNDW | 0.86 | -0.05 | 0.21 | -0.26 | 0.87 | CNDW | 0.14 | 0.58 | 0.26 | 0.43 |
| | 0.93 | -0.15 | 0.53 | -0.64 | | | 0.12 | 0.62 | 0.38 | |
| NSMIL | 0.77 | -0.36 | 0.26 | -0.02 | 0.80 | NSMIL | -0.57 | 0.25 | 0.09 | 0.43 |
| | 0.87 | -0.63 | 0.51 | -0.13 | | | -0.62 | 0.39 | 0.12 | |
| DDL | 0.72 | -0.42 | 0.01 | 0.31 | 0.78 | DDL | -0.53 | -0.11 | -0.14 | 0.32 |
| | 0.83 | -0.66 | 0.01 | 0.50 | | | -0.54 | -0.08 | -0.20 | |
| YW | 0.16 | 0.67 | 0.65 | 0.19 | 0.95 | YW | -0.30 | -0.57 | 0.61 | 0.70 |
| | 0.56 | 0.95 | 0.95 | 0.65 | | | -0.38 | -0.67 | 0.72 | |
| TAX, % | 83.7 | 25.7 | 22.9 | 43.4 | | | 80.4 | 56.4 | 42.8 | |
| SD, % | 9.4 | 4.5 | 25.5 | 2.9 | | | 3.5 | 14.7 | 2.27 | |

$r^2$ – squared multiple correlation for variables/indices according to multivariate regression. MSL – maximal skull length, NSMIL – nasal bone minimal length, FCL – facial length, PL – palatal length, RSL – rostral length, DUL – length of upper diastema, TUL – alveolar length of upper tooth-row, FGL – length of *fossa glenoidea* (glenoid cavity) (*fg*), ZW – zygomatic width, YW – interorbital width, MSW – mastoid width, RSMIW, RSMEW, RSMAW – minimal, intermediate and maximal width of rostrum, INCW – double upper incisive width, FGMIW, FGMEW, FGMAW – minimal, intermediate and maximal width of skull base between left and right glenoid *fossa glenoidea*, RSMIH, RSMEH, RSMAH – minimal, intermediate and maximal height of rostrum, CPSH – skull height, MNDL – mandible length, DDL – length of lower diastema, TDL – alveolar length of lower tooth-row, MNDH – height of horizontal branch of mandible, and, in addition, CNDL, CNDW – the length and width of condyle.

**Table S4** Pearson correlations of morphometric variables (VAR) and indices (VAR/ML) with Principal Component (PC) coordinates PC1–PC4 based on the coordinates of the SZM and SHM morphospace models; statistically significant $r$ values ($p < 0.001$) are highlighted in red

| VAR | PC coordinate | | | |
|---|---|---|---|---|
| VAR/MSL | PC1 | PC2 | PC3 | PC4 |
| MSL | 0.93 | -0.18 | -0.38 | 0.22 |
| RSMIH | 0.95 | -0.09 | -0.33 | 0.18 |
| RSMIH/ML | 0.83 | 0.03 | -0.22 | 0.09 |
| MSW | 0.94 | -0.10 | -0.29 | 0.24 |
| MSW/ML | 0.02 | 0.27 | 0.30 | 0.05 |
| ZW | 0.92 | -0.12 | -0.42 | 0.21 |
| ZW/ML | 0.57 | 0.09 | -0.38 | 0.09 |
| RSMEW | 0.97 | -0.16 | -0.19 | 0.17 |
| RSMEW/ML | 0.84 | -0.13 | 0.09 | 0.07 |
| RSMAW | 0.96 | -0.10 | -0.25 | 0.18 |
| RSMAW/ML | 0.69 | 0.09 | 0.06 | 0.07 |
| PL | 0.91 | -0.17 | -0.42 | 0.22 |
| PL/ML | -0.04 | 0.02 | -0.22 | 0.00 |
| FGMEW | 0.95 | -0.19 | -0.18 | 0.25 |
| FGMEW/ML | 0.10 | -0.04 | 0.54 | 0.08 |
| INCW | 0.96 | -0.24 | -0.19 | 0.11 |
| INCW/ML | 0.85 | -0.28 | 0.05 | -0.02 |
| FGMAW | 0.94 | -0.23 | -0.16 | 0.26 |
| FGMAW/ML | 0.10 | -0.14 | 0.57 | 0.11 |
| MNDL | 0.89 | 0.00 | -0.43 | 0.25 |
| MNDL/ML | -0.10 | 0.51 | -0.13 | 0.07 |
| DUL | 0.85 | -0.17 | -0.55 | 0.18 |
| DUL/ML | 0.02 | -0.05 | -0.65 | -0.08 |
| FGMIW | 0.92 | -0.08 | -0.20 | 0.26 |
| FGMIW/ML | 0.09 | 0.21 | 0.37 | 0.13 |
| FCL | 0.82 | -0.19 | -0.51 | 0.32 |
| FCL/ML | 0.07 | -0.11 | -0.47 | 0.37 |
| RSL | 0.84 | -0.27 | -0.33 | 0.31 |
| RSL/ML | 0.14 | -0.28 | -0.01 | 0.29 |
| CPSH | 0.82 | -0.14 | -0.47 | 0.29 |
| CPSH/ML | -0.12 | 0.05 | -0.25 | 0.21 |
| MNDH | 0.84 | 0.06 | -0.46 | 0.21 |
| MNDH/ML | 0.35 | 0.41 | -0.38 | 0.10 |
| CNDL | 0.86 | -0.20 | -0.29 | 0.29 |
| CNDL/ML | 0.24 | -0.11 | 0.03 | 0.26 |
| RSMAH | 0.84 | 0.06 | -0.35 | 0.29 |
| RSMAH/ML | 0.14 | 0.41 | -0.07 | 0.22 |
| RSMIW | 0.84 | 0.15 | -0.38 | 0.24 |
| RSMIW/ML | 0.37 | 0.53 | -0.20 | 0.16 |
| RSMEH | 0.82 | -0.17 | -0.40 | 0.27 |
| RSMEH/ML | 0.27 | -0.09 | -0.21 | 0.20 |
| TUL | 0.95 | -0.16 | 0.04 | 0.13 |
| TUL/ML | 0.54 | -0.07 | 0.55 | -0.05 |
| FGL | 0.86 | -0.42 | -0.22 | 0.14 |
| FGL/ML | 0.21 | -0.59 | 0.16 | -0.08 |
| TDL | 0.92 | -0.13 | 0.07 | 0.21 |
| TDL/ML | 0.45 | -0.01 | 0.62 | 0.09 |
| CNDW | 0.83 | 0.19 | -0.31 | 0.26 |
| CNDW/ML | 0.34 | 0.56 | -0.11 | 0.18 |
| NSMIL | 0.66 | 0.12 | -0.62 | 0.24 |
| NSMIL/ML | -0.35 | 0.44 | -0.39 | 0.04 |

| VAR<br>VAR/MSL | PC coordinate | | | |
|---|---|---|---|---|
| | PC1 | PC2 | PC3 | PC4 |
| DDL | 0.61 | -0.18 | -0.61 | 0.10 |
| DDL/ML | -0.47 | -0.00 | -0.28 | -0.18 |
| YW | 0.12 | -0.10 | 0.31 | 0.72 |
| YW/ML | -0.64 | 0.08 | 0.53 | 0.33 |

MSL – maximal skull length, NSMIL – minimal nasal bone length, FCL – facial length, PL – palatal length, RSL – rostral length, DUL – length of upper diastema, TUL – alveolar length of upper tooth-row, FGL – length of *fossa glenoidea* (glenoid cavity) (*fg*), ZW – zygomatic width, YW – interorbital width, MSW – mastoid width, RSMIW, RSMEW, RSMAW – minimal, intermediate and maximal width of rostrum, INCW – double upper incisive width, FGMIW, FGMEW, FGMAW – minimal, intermediate and maximal width of skull base between left and right glenoid *fossa glenoidea*, RSMIH, RSMEH, RSMAH – minimal, intermediate and maximal height of rostrum, CPSH – skull height, MNDL – mandible length, DDL – length of lower diastema, TDL – alveolar length of lower tooth-row, MNDH – height of horizontal branch of mandible, and, in addition, CNDL, CNDW – the length and width of condyle.

**Supplementary materials for the section *A morphological comparison of the North Caucasian Spalax sp. (2n = 62) with members of the genus Spalax***

The North Caucasian blind mole rat is a typical member of the genus *Spalax* based on a combination of the following craniometric characteristics (Topachevski 1969).

The topography of the mandibular fossa is important for the temporal and spatial separation of the main functions of the mandibular apparatus. When chewing, the head of condyloid process is in the posterior third of the *fossa glenoidea*, when chewing - in the middle, when biting - it moves within the middle third of the fossa; when sharpening the lower incisors, it occupies the most anterior position in the fossa.

Before biting, the mole rat opens its mouth wide. The maximum-recorded mouth-opening angle is 46°. The condyle of the condyloid process is within the middle third of the *fossa glenoidea*. The lower incisors are spread apart; the distance between them can be up to 6.8 mm. The first phase of biting is the forward and upward movement of the lower incisors, with them sinking into the clay for a distance equal to their free length. The second phase is the upward or up-backward movement of the incisors towards the upper incisors, which act as a support. After the lower and upper incisors have closed, the condyle of the articular process moves to the posterior third of the *fossa glenoidea*. As the mouth closes, the medial surfaces of the lower incisors converge.

In *Spalax* sp. (2*n* = 62), short glenoid cavities combined with the relatively small difference in distance between their anterior and posterior edges result in a rather sharp angle between the imaginary lines drawn along the surfaces of the glenoid cavities (Fig. S8A, Supporting information). This divergence of the incisors is the result of pressure from *fossa glenoidea* outer edges of fossa on the head of the condylar process. A potentially distance between the incisor tips depends on an angle between the *fossa glenoidea*. The incisors will more widely spread if this angle will more and, therefore, a size of piece of soil that the animal can bite off (in a single digging act) will also dependent on this angle. The biological interpretation of this result, if of course it is confirmed by more accurate measurements, could be a) the North Caucasian blind mole rat is more adapted to digging in dense soils, or b) it generally has a relatively low adaptation to burrowing among all other *Spalax*. We cannot exclude a compensation for the described feature of a base of skull in the North Caucasian blind mole rat by the relatively long condylar process

**Table S5** Statistical comparison (Welch test = unequal variances *t*-test (Welch 1947)) of the means of skull measurements in *Spalax* sp. (2n = 62) and all the other species of genus *Spalax* except *S. zemni*

| VAR<br>VAR/ML, % | ***Spalax* sp., *N*=18** | ***S. graecus*, *N*=21** | **Welch F, *p*** | ***S. microphthalmus*, *N*=43** | **Welch F, *p*** | ***S. arenarius*, *N*=19** | **Welch F, *p*** | ***S. giganteus*, *N*=25** | **Welch F, p** | ***S. uralensis*, *N*=13** | **Welch F, *p*** |
|---|---|---|---|---|---|---|---|---|---|---|---|
| MSL | 50.4±0.47 | 55.5±0.60 | 44.9, <0.0001 | 55.0±0.42 | 52.0, <0.0001 | 53.9±0.48 | 26.2.0, <0.0001 | 67.9±1.0 | 247.2, <0.0001 | 61.3±1.12 | 79.6, <0.0001 |
| MSW | 26.3±0.23 | 27.6±0.27 | 13.9, 0.0006 | 27.3±0.22 | 9.2, 0.004 | 26.8±0.28 | 2.1, n.s. | 34.2±0.28 | 200.7, <0.0001 | 30.9±0.51 | 68.3, <0.0001 |
| MSW% | 52.2±0.32 | 50.0±0.45 | 16.1, 0.0003 | 49.6±0.24 | 41.9, <0.0001 | 49.8±0.4 | 21.4, 0.0001 | 50.4±0.4 | 17.7, 0.0002 | 50.5±0.42 | 9.9, 0.0044 |
| ZW | 37.5±0.53 | 41.6±0.68 | 23, <0.0001 | 41.8±0.43 | 39.4, <0.0001 | 39.8±0.47 | 11.1, 0.0021 | 53.1±0.47 | 186.2, <0.0001 | 48.5±0.98 | 98.3, <0.0001 |
| ZW% | 74.3±0.56 | 75.2±0.71 | 0.9, n.s. | 75.9±0.34 | 5.6, 0.0243 | 73.9±0.47 | 0.3, n.s | 78.2±0.47 | 23.8, <0.0001 | 79.2±0.45 | 45.3, <0.0001 |
| RSMIW | 8.5±0.21 | 9.0±0.15 | 3.2, n.s. | 8.95±0.13 | 3.4, n.s. | 7.81±0.13 | 7.8, 0.0091 | 11.5±0.13 | 75.2, <0.0001 | 10.8±0.36 | 30.1, <0.0001 |
| RSMIW% | 16.9±0.35 | 16.2±0.27 | 2.0, n.s. | 16.3±0.15 | 2.4, n,s, | 14.5±0.21 | 33, <0.0001 | 17±0.21 | 0.1, n.s. | 17.5±0.39 | 1.7, n.s. |
| RSMEW | 11±0.14 | 12.2±0.17 | 26.5, <0.0001 | 11.8±0.12 | 18.7, 0.0001 | 12.3±0.12 | 48.6, <0.0001 | 17.4±0.12 | 287.5, <0.0001 | 16.4±0.28 | 300.8, <0.0001 |
| RSMEW% | 21.9±0.19 | 22±0.21 | 0.3, n.s. | 21.5±0.11 | 3.3, n.s. | 22.9±0.18 | 15.6, 0.0004 | 25.5±0.18 | 174.4, <0.0001 | 26.8±0.17 | 385.9, <0.0001 |
| RSMAW | 11.7±0.2 | 12.6±0.14 | 15.9, 0.0004 | 12.5±0.15 | 11.5, 0.0016 | 12.3±0.13 | 8.4, 0.0071 | 16.9±0.13 | 240.6, <0.0001 | 15.9±0.27 | 159.9, <0.0001 |
| RSMAW% | 23.1±0.29 | 22.9±0.23 | 0.5, n.s. | 22.7±0.14 | 1.5, n.s. | 22.9±0.19 | 0.3, n.s. | 24.9±0.19 | 27.1, <0.0001 | 25.9±0.24 | 57.3, <0.0001 |
| RSMIH | 6.6±0.09 | 6.9±0.12 | 2.2, n.s. | 7.11±0.09 | 13.8, 0.0005 | 7.09±0.09 | 13.4, 0.0008 | 10.5±0.09 | 190.7, <0.0001 | 9±0.23 | 90.2, <0.0001 |
| RSMIH% | 13.2±0.09 | 12.4±0.14 | 20.5, 0.0001 | 12.9±0.1 | 3.3, n,s. | 13.2±0.12 | 0.0, n.s. | 15.4±0.12 | 91.1, <0.0001 | 14.7±0.19 | 50.5, <0.0001 |
| RSMEH | 7.9±0.12 | 8.9±0.17 | 24.9, <0.0001 | 9.02±0.12 | 47.8, <0.0001 | 8.7±0.14 | 20, 0.0001 | 11.6±0.14 | 141.2, <0.0001 | 9.73±0.22 | 54.5, <0.0001 |
| RSMEH% | 15.6±0.23 | 16.1±0.3 | 1.7, n.s. | 16.4±0.16 | 8.1, 0.0075 | 16.2±0.26 | 2.6, n.s. | 17.1±0.26 | 15.2, 0.0003 | 15.9±0.2 | 0.8, n.s. |
| RSMAH | 12.9±0.21 | 13.4±0.23 | 3.1, n.s. | 12.8±0.21 | 0.2, n.s. | 12.9±0.17 | 0.0, n.s. | 16.3±0.17 | 76, <0.0001 | 15.8±0.45 | 34.3, <0.0001 |
| RSMAH% | 25.6±0.35 | 24.3±0.28 | 8.7, 0.0056 | 23.2±0.29 | 27.7, <0.0001 | 24.0±0.15 | 18.1, 0.0003 | 24.0±0.15 | 14.2, 0.0007 | 25.7±0.37 | 0.1, n.s. |
| NSMIL | 19.6±0.34 | 20.3±0.37 | 1.8, n.s. | 21.1±0.26 | 13.1, 0.0009 | 19.3±0.43 | 0.2, n.s. | 23.6±0.43 | 38.6, <0.0001 | 21.8±0.64 | 9.2, 0.007 |
| NSMIL% | 38.8±0.49 | 36.6±0.47 | 10.7, 0.0024 | 38.4±0.34 | 0.4, n.s. | 35.9±0.69 | 12, 0.0015 | 34.7±0.69 | 37.1, <0.0001 | 35.5±0.69 | 15.4, 0.0007 |
| RSL | 16.7±0.28 | 17.9±0.24 | 10.8, 0.0023 | 17.1±0.19 | 1.2, n.s. | 18.6±0.29 | 21.3, 0.0001 | 22.8±0.29 | 107.5, <0.0001 | 19.6±0.4 | 35.3, <0.0001 |
| RSL% | 33.1±0.35 | 32.4±0.36 | 2.1, n.s. | 31.1±0.23 | 24.4, <0.0001 | 34.5±0.42 | 6.0, 0.0197 | 33.5±0.42 | 0.4, n.s. | 32.0±0.35 | 5.3, 0.0296 |
| FCL | 28.5±0.59 | 31.8±0.48 | 18.9, 0.0001 | 30.8±0.36 | 11.4, 0.002 | 31.1±0.44 | 12.7, 0.0012 | 39.2±0.44 | 97.1, <0.0001 | 32.7±0.69 | 21.7, 0.0001 |
| FCL% | 56.4±0.81 | 57.5±0.45 | 1.2, n.s. | 56±0.36 | 0.2, n.s. | 57.8±0.55 | 1.8, n.s. | 57.6±0.55 | 1.4, n.s. | 53.4±0.42 | 11.1, 0.0027 |
| YW | 7.7±0.12 | 8.2±0.14 | 7.2, 0.0109 | 7.43±0.12 | 3.1, n.s. | 7.66±0.09 | 0.2, n.s. | 8.02±0.09 | 1.9, n.s. | 7.61±0.17 | 0.3, n.s. |
| YW% | 15.4±0.28 | 14.9±0.31 | 1.2, n.s. | 13.5±0.24 | 24.4, <0.0001 | 14.3±0.24 | 8.9, 0.0052 | 11.9±0.24 | 74.3, <0.0001 | 12.5±0.38 | 36.6, <0.0001 |
| PL | 27.1±0.27 | 29.1±0.35 | 20.8, 0.0001 | 29.5±0.26 | 40.2, <0.0001 | 29.2±0.35 | 21.5, 0.0001 | 36.3±0.35 | 208.5, <0.0001 | 32.6±0.59 | 71.8, <0.0001 |
| PL% | 53.8±0.24 | 52.7±0.21 | 13.1, 0.0009 | 53.6±0.21 | 0.6, n.s. | 54.2±0.35 | 0.6, n.s. | 53.4±0.35 | 1.4, n.s. | 53.3±0.3 | 2.0, n.s. |
| DUL | 18.7±0.31 | 20.1±0.32 | 9.9, 0.0033 | 21.1±0.21 | 43.4, <0.0001 | 20.5±0.25 | 21, 0.0001 | 25.6±0.25 | 138.2, <0.0001 | 22.8±0.43 | 60.1, <0.0001 |
| DUL% | 37.0±0.34 | 36.2±0.35 | 2.2, n.s. | 38.4±0.16 | 14.9, 0.0007 | 38±0.26 | 6.2, 0.0181 | 37.6±0.26 | 1.9, n.s. | 37.2±0.36 | 0.3, n.s. |
| CPSH | 21.2±0.3 | 23.9±0.3 | 43.4, <0.0001 | 23.7±0.26 | 41.1, <0.0001 | 23.4±0.25 | 34.7, <0.0001 | 28.5±0.25 | 110.1, <0.0001 | 25.7±0.61 | 45.0, <0.0001 |
| CPSH% | 41.9±0.3 | 43.3±0.46 | 6.1, 0.0187 | 43.0±0.25 | 8, 0.0075 | 43.5±0.43 | 9.1, 0.0049 | 41.9±0.43 | 0.0, n.s. | 41.9±0.45 | 0.0, n.s. |

| VAR<br>VAR/ML, % | ***Spalax* sp., *N*=18** | ***S. graecus, N*=21** | **Welch F, *p*** | ***S. microphthalmus, N*=43** | **Welch F, *p*** | ***S. arenarius, N*=19** | **Welch F, *p*** | ***S. giganteus, N*=25** | **Welch F, p** | ***S. uralensis, N*=13** | **Welch F, *p*** |
|---|---|---|---|---|---|---|---|---|---|---|---|
| TUL | 7.70±0.09 | 8.50±0.1 | 41.8, <0.0001 | 8.28±0.08 | 25.7, <0.0001 | 8.51±0.09 | 41.8, <0.0001 | 11.6±0.09 | 589.2, <0.0001 | 11±0.22 | 192.8, <0.0001 |
| TUL% | 15.2±0.25 | 15.5±0.24 | 0.5, n.s, | 15.1±0.17 | 0.3, n.s. | 15.8±0.17 | 3.5, n.s. | 17.1±0.17 | 29.6, <0.0001 | 17.9±0.27 | 54.3, <0.0001 |
| INCW | 7.5±0.1 | 8.6±0.09 | 59.9, <0.0001 | 8.41±0.07 | 54.3, <0.0001 | 8.87±0.1 | 95.7, <0.0001 | 13.1±0.1 | 388.9, <0.0001 | 11.5±0.29 | 169.4, <0.0001 |
| INCW% | 14.8±0.15 | 15.5±0.15 | 9.2, 0.0045 | 15.3±0.09 | 6.9, 0.0132 | 16.5±0.16 | 54.6, <0.0001 | 19.3±0.16 | 253.4, <0.0001 | 18.9±0.38 | 95.5, <0.0001 |
| FGMIW | 18.7±0.26 | 19.5±0.17 | 6.3, 0.0177 | 19.1±0.12 | 1.9, n.s. | 18.8±0.2 | 0.0, n.s. | 24.6±0.2 | 169.1, <0.0001 | 22.1±0.39 | 50.6, <0.0001 |
| FGMIW% | 37.1±0.42 | 35.3±0.31 | 12.4, 0.0013 | 34.8±0.21 | 24.3, <0.0001 | 34.8±0.39 | 16.0, 0.0003 | 36.2±0.39 | 2.7, n.s. | 36.1±0.48 | 2.9, n.s. |
| FGMEW | 20.6±0.19 | 22.3±0.18 | 42.2, <0.0001 | 21.8±0.13 | 24.3, <0.0001 | 21.8±0.21 | 17.8, 0.0002 | 27.8±0.21 | 325, <0.0001 | 25±0.36 | 117.3, <0.0001 |
| FGMEW% | 40.9±0.27 | 40.4±0.34 | 1.5, n.s. | 39.6±0.23 | 14.2, 0.0005 | 40.6±0.52 | 0.4, n.s. | 41±0.52 | 0.0, n.s. | 40.8±0.35 | 0.0, n.s. |
| FGMAW | 21.7±0.23 | 23.6±0.18 | 43.0, <0.0001 | 23.0±0.13 | 22.9, <0.0001 | 22.9±0.19 | 16.9, 0.0002 | 29.5±0.19 | 368.6, <0.0001 | 26.4±0.28 | 173.3, <0.0001 |
| FGMAW% | 43.1±0.34 | 42.8±0.43 | 0.4, n.s. | 41.8±0.26 | 9.2, 0.0044 | 42.6±0.36 | 1.0, n.s. | 43.6±0.36 | 1.0, n.s. | 43.2±0.51 | 0, n.s. |
| FGL | 9.3±0.2 | 10.7±0.16 | 31.1, <0.0001 | 10.7±0.12 | 37.8, <0.0001 | 11.5±0.14 | 85.1, <0.0001 | 13.7±0.14 | 240.8, <0.0001 | 12.8±0.22 | 134.8, <0.0001 |
| FGL% | 18.4±0.29 | 19.4±0.26 | 6.8, 0.0129 | 19.5±0.16 | 11.7, 0.0019 | 21.4±0.26 | 62.1, <0.0001 | 20.2±0.26 | 29.2, <0.0001 | 20.9±0.29 | 37.9, <0.0001 |
| MNDH | 8.8±0.12 | 9.3±0.15 | 6.8, 0.013 | 9.95±0.16 | 34.8, <0.0001 | 8.74±0.13 | 0,0, n.s. | 12.4±0.13 | 127.4, <0.0001 | 11±0.25 | 62.0, <0.0001 |
| MDH/<br>MNDL, % | 27.9±0.33 | 28.3±0.32 | 0.9, n.s. | 29.7±0.3 | 16.3, 0.0002 | 27.3±0.42 | 1.4, n.s. | 30.4±0.42 | 21.8, <0.0001 | 29.6±0.39 | 11.6, 0.0022 |
| MDH% | 17.4±0.14 | 16.8±0.21 | 5.8, 0.0214 | 18.1±0.2 | 8.3, 0.0055 | 16.2±0.22 | 19.4, 0.0001 | 18.2±0.22 | 8.1, 0.0073 | 17.9±0.25 | 3.8, n.s. |
| MNDL | 31.4±0.37 | 32.7±0.31 | 7.5, 0.0095 | 33.5±0.4 | 14.9, 0.0003 | 32±0.29 | 1.9, n.s. | 40.6±0.29 | 155, <0.0001 | 37±0.65 | 56.7, <0.0001 |
| MNDL/MSL% | 62.3±0.44 | 59.1±0.32 | 32.2, <0.0001 | 60.8±0.47 | 4.9, 0.0316 | 59.5±0.31 | 26.6, <0.0001 | 59.9±0.31 | 14.9, 0.0004 | 60.4±0.49 | 7.6, 0.0103 |
| DDL | 7.9±0.14 | 8.3±0.13 | 3.4, 0.0722 | 9.02±0.11 | 38, <0.0001 | 8.79±0.14 | 19, 0.0001 | 9.92±0.14 | 61.6, <0.0001 | 9.03±0.17 | 25, <0.0001 |
| DDL% | 15.7±0.19 | 15.0±0.24 | 5.3, 0.0267 | 16.4±0.14 | 8.5, 0.006 | 16.3±0.22 | 4.5, 0.0416 | 14.6±0.22 | 15.7, 0.0003 | 14.8±0.27 | 8.0, 0.0093 |
| TDL | 7.1±0.07 | 8.0±0.09 | 68.2, <0.0001 | 7.49±0.05 | 23.3, <0.0001 | 7.78±0.1 | 34.7, <0.0001 | 10.5±0.1 | 534.5, <0.0001 | 9.6±0.15 | 239.8, <0.0001 |
| TDL% | 14.1±0.25 | 14.5±0.2 | 1.9, n.s. | 13.7±0.12 | 2.2, n.s. | 14.5±0.21 | 1.5, n.s. | 15.6±0.21 | 17.7, 0.0001 | 15.7±0.24 | 21.9, 0.0001 |
| CONDL | 5±0.12 | 5.4±0.07 | 7.0, 0.0132 | 5.2±0.06 | 2.1, n.s. | 5.39±0.07 | 7.7, 0.0101 | 6.9±0.07 | 115.6, <0.0001 | 6.03±0.14 | 29.1, <0.0001 |
| CONDL% | 9.9±0.18 | 9.7±0.1 | 0.6, n.s. | 9.46±0.07 | 5, 0.0348 | 10.0±0.09 | 0.3, n.s. | 10.2±0.09 | 1.5, n.s. | 9.85±0.22 | 0.0, n.s. |
| CONDW | 2.8±0.05 | 2.9±0.08 | 2.0, n.s. | 2.93±0.04 | 4.4, 0.043 | 2.53±0.04 | 17.5, 0.0002 | 3.84±0.04 | 107.5, <0.0001 | 3.42±0.11 | 24.6, 0.0001 |
| CONDW% | 5.6±0.07 | 5.3±0.11 | 4.3, 0.0445 | 5.33±0.06 | 5.7, 0.0221 | 4.7±0.05 | 88.9, <0.0001 | 5.7±0.05 | 0.7, n.s. | 5.6±0.15 | 0.0, n.s. |

MSL – maximal skull length, NSMIL – minimal nasal bone length, FCL – facial length, PL – palatal length, RSL – rostral length, DUL – length of upper diastema, TUL – alveolar length of upper tooth-row, FGL – length of *fossa glenoidea* (glenoid cavity) (*fg*), ZW – zygomatic width, YW – interorbital width, MSW – mastoid width, RSMIW, RSMEW, RSMAW – minimal, intermediate and maximal width of rostrum, INCW – double upper incisive width, FGMIW, FGMEW, FGMAW – minimal, intermediate and maximal width of skull base between left and right glenoid *fossa glenoidea*, RSMIH, RSMEH, RSMAH – minimal, intermediate and maximal height of rostrum, CPSH – skull height, MNDL – mandible length, DDL – length of lower diastema, TDL – alveolar length of lower tooth-row, MNDH – height of horizontal branch of mandible, and, in addition, CNDL, CNDW – the length and width of condyle.

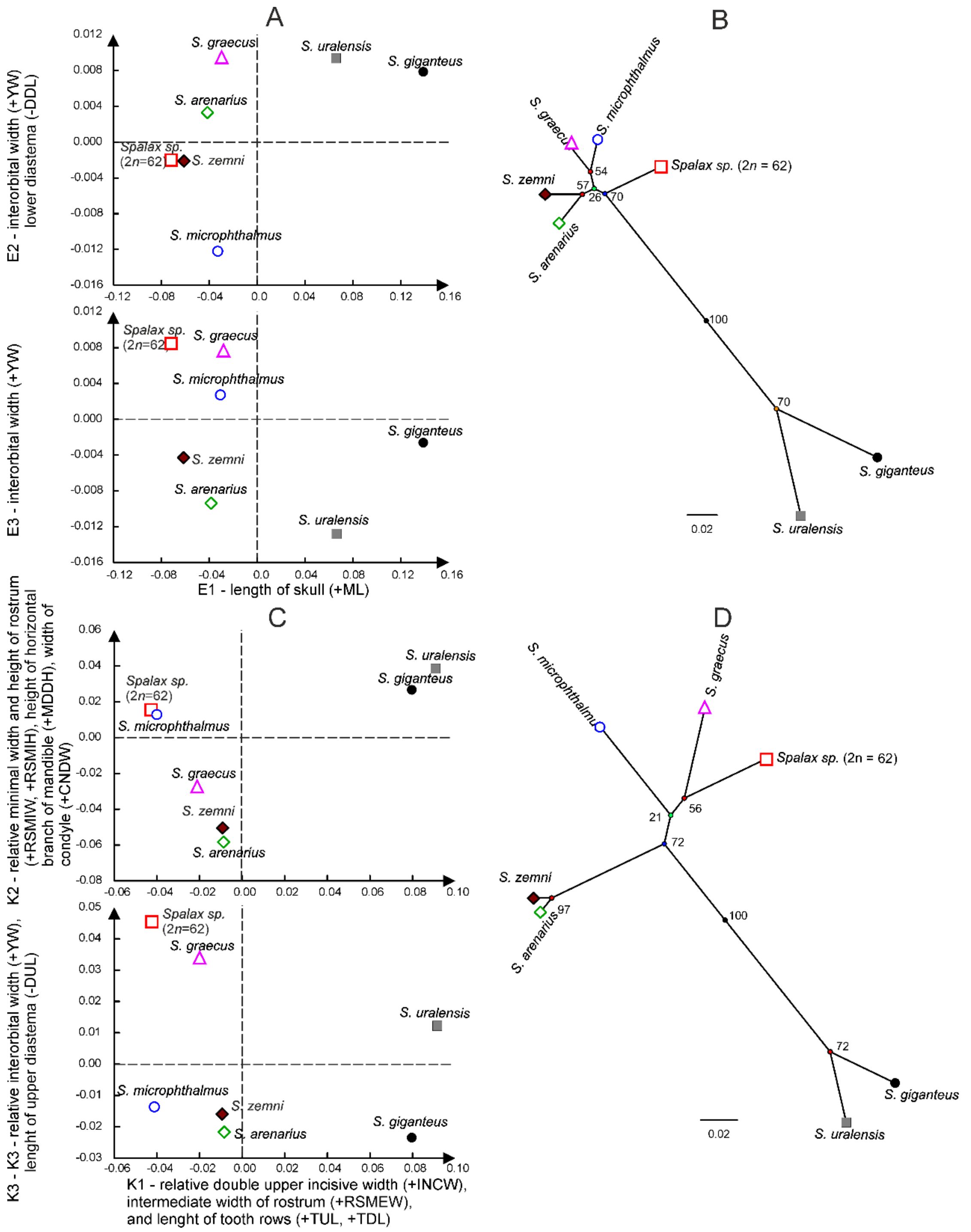


**Figure S3** The projections of the *Spalax* species sample centroids onto the E1xE2, E1xE3 (A), K1xK2, and K1xK3 (C) coordinates of the SZM and SHM models, respectively, and the radial classification trees (Euclid distances, UPGMA method) of the species centroids based on the E1 – E4 coordinates of the SZM model (B) and K1 – K3 coordinates of the SHM model (D). The numbers near the nodes of the trees estimate bootstrap support as a percentage of 1000 iterations.

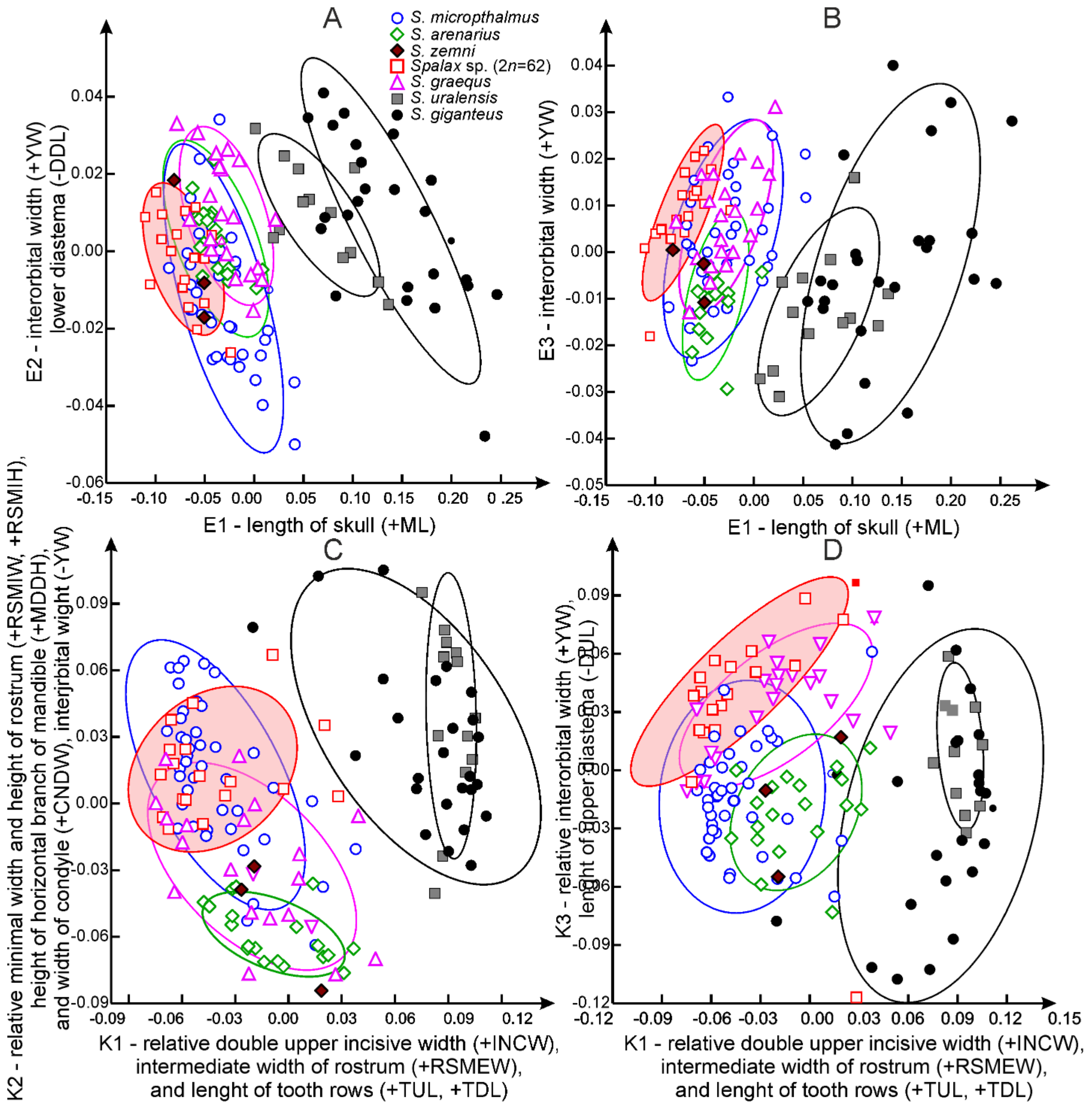


**Figure S4** The projections of the species of the *Spalax* samples (specimens) onto the E1x E2 (A), E1xE3 (B), K1x K2 (C) and K1xK3 (D) coordinates of the SZM (A, B) and SHM (C, D) models.

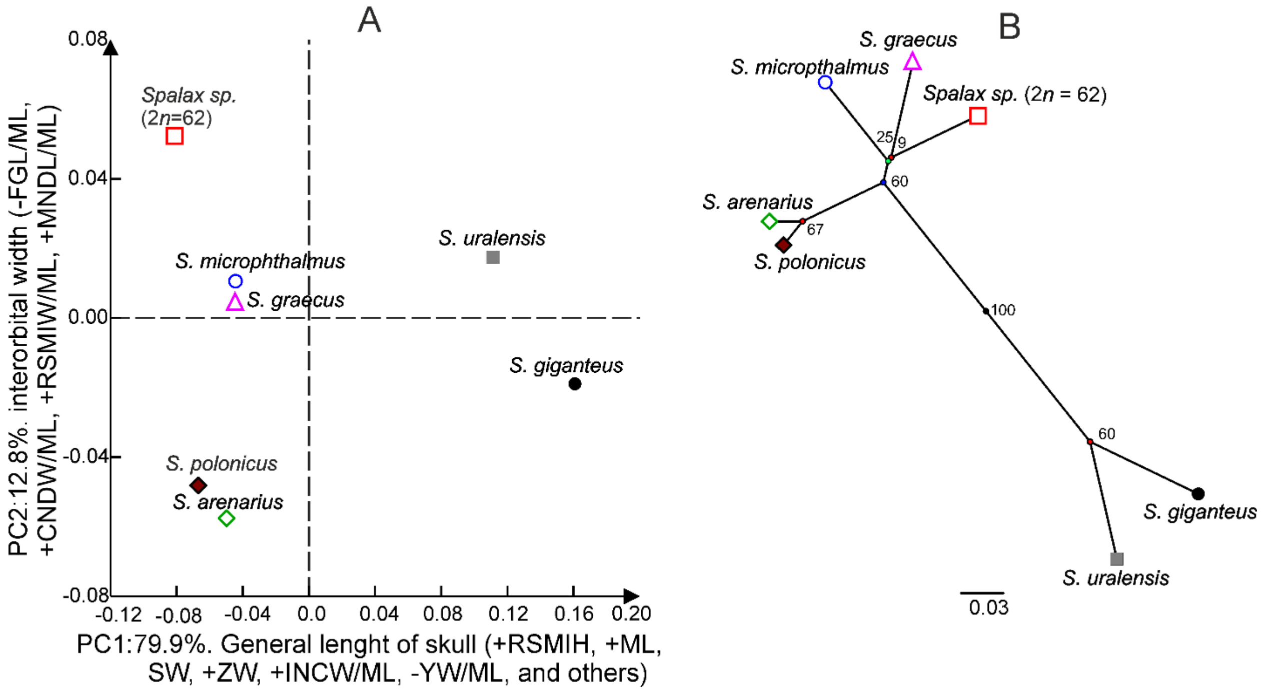


**Figure S5** The projection of the *Spalax* species sample centroids onto PC1 and PC2 (A) based on the E1 – E4 coordinates of the SZM model and K1 – K3 coordinates of the SHM model, and the radial classification tree (Euclid distances, UPGMA method) of the species centroids based on PC1-PC4 that accounted for 99.8 % of the E1-E4 and K1-K3 variance.

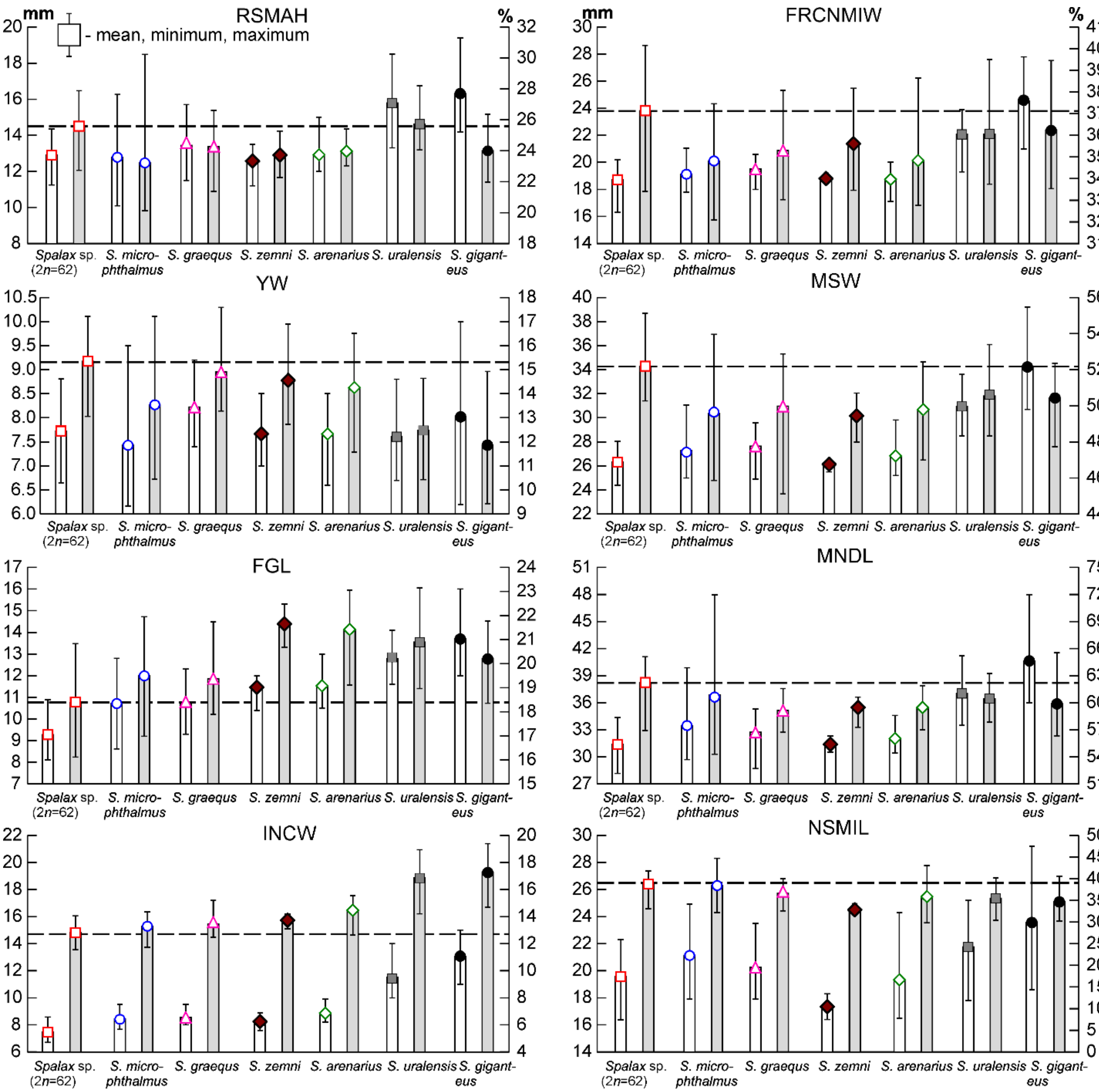


**Figure S6** Absolute (white rectangles) values of some skull measurements and their indices (grey rectangles) in members of the genus *Spalax*.

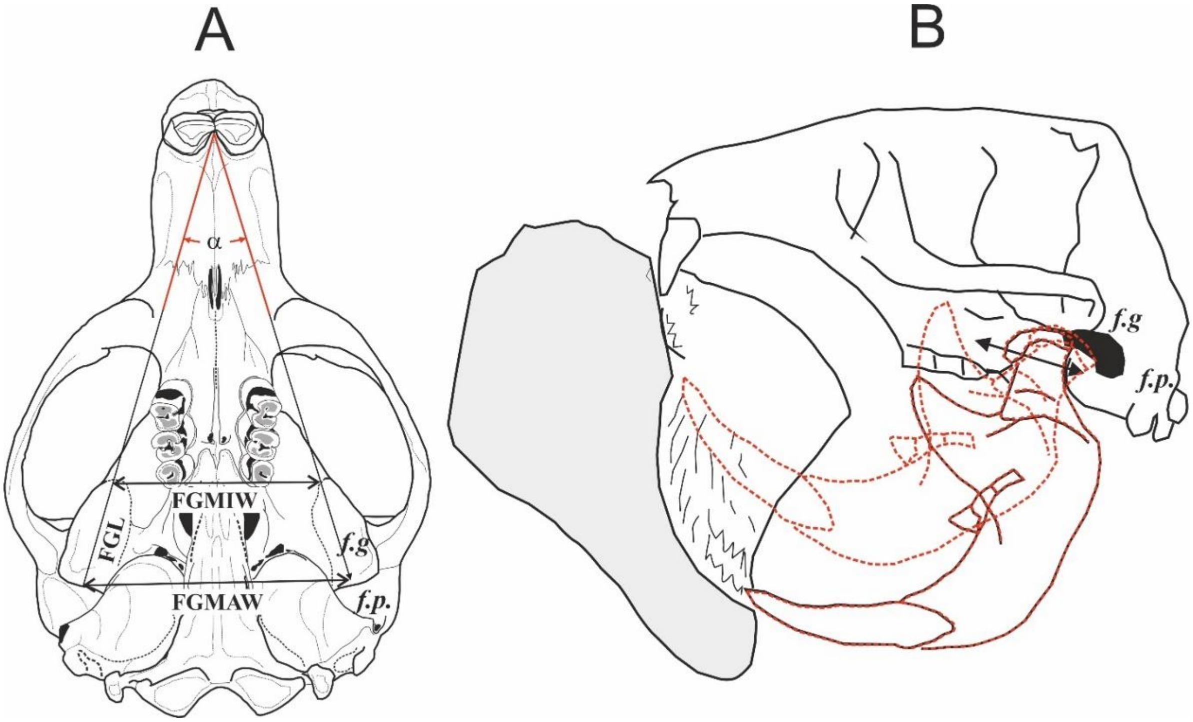


**Figure S7** A - Imaginary angle (α) between glenoid cavities; FGL – length of *fossa glenoidea*, FGMIW, FGMAW – minimal and maximal width of skull base between the left and right *fossa glenoidea*. B - Schematic of the position of the mandible in the initial and intermediate (red dotted line) phases of the digging according to (Zubtsova 1986) with additions; *f.g.* - *fossa glenoidea*, *f.p* - *fossa pseudoglenoidea*.

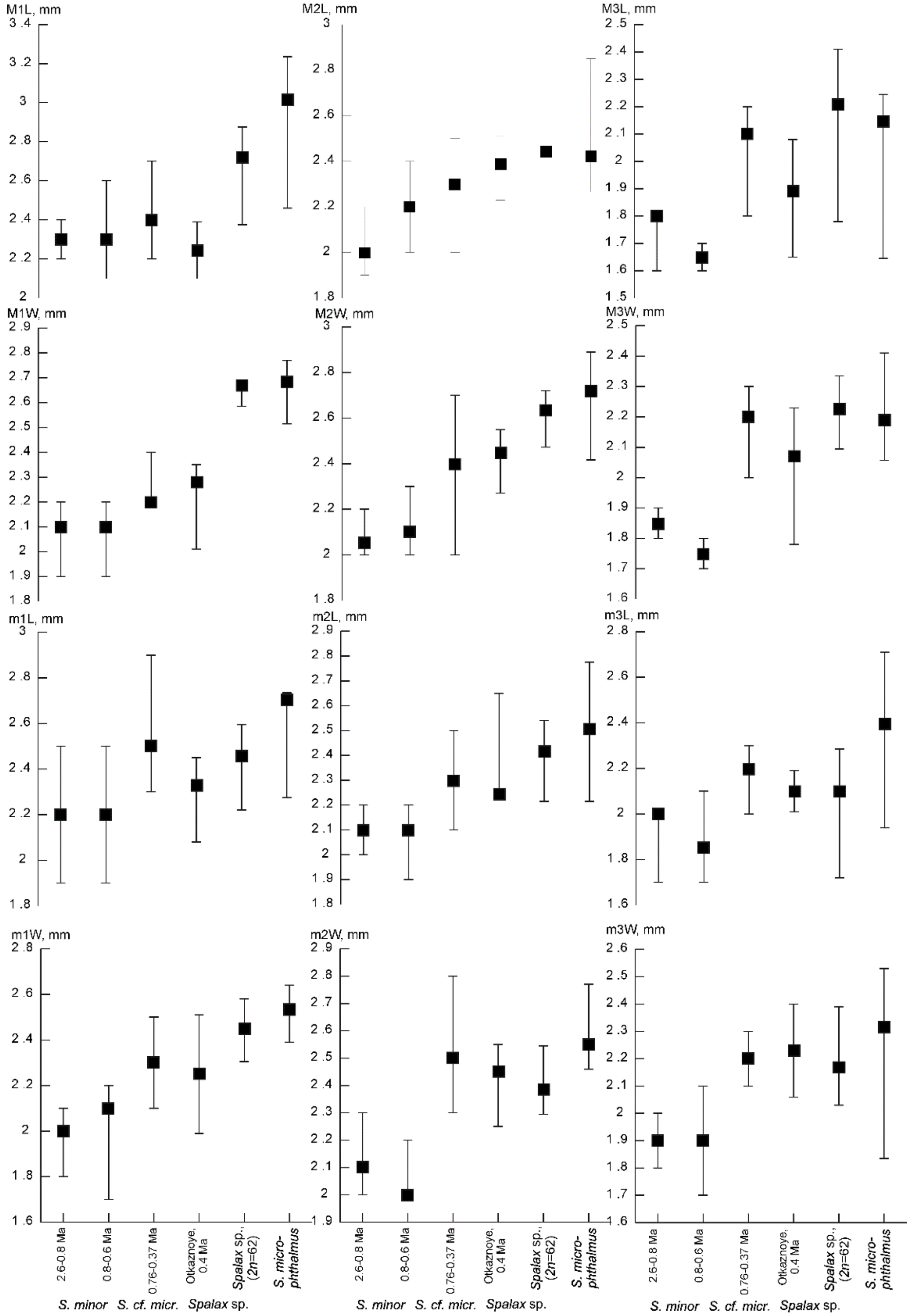


**Figure S8** The medians and the minimum-maximum values of the length (L) and width (W) of the upper (M1-M3) and lower (m1-m3) molars of *S. minor* from the Early Pleistocene – the beginning of the Middle Pleistocene from the lower Dnieper and the Azov-Black Sea region, and *S*. cf. *microphthalmus* (Stadnik 2009), *Spalax* sp. from the Otkaznoye locality - middle of the Middle Pleistocene (material collected and studied by Dr. A.K. Markova); and *Spalax* sp. (2n = 62) (Kabardino-Balkaria, Russia) and *S. microphthalmus* (mainly a population sample from the Kursk district, Russia).
